# Spatial confinement of gene drives: Assessing risk of failure using global sensitivity analysis

**DOI:** 10.64898/2026.02.18.706608

**Authors:** Cole D. Butler, Alun L. Lloyd

## Abstract

Gene drives allow pest populations to be genetically modified to reduce their harm on agriculture and human health. The genetic modification, or payload, spreads within a target population at rates exceeding normal Mendelian inheritance. While gene drives have demonstrated immense potential in laboratory populations, they present unique challenges. Foremost among these challenges is spatial confinement, or ensuring that the payload remains confined to target populations. However, there is an inherent tension between gene drive spread and spatial confinement: increasing the spreading efficiency of a gene drive increases the risk of escape, while engineering confinement mechanisms increases the risk of gene drive extinction. In this work, we explore spatial outcomes in gene drives designed for spatial confinement and the dependence of these outcomes on target organism dispersal and payload fitness cost. We use a stochastic spatial model to compute the probability of failure for each gene drive, and use techniques from global sensitivity analysis to quantify the contribution of dispersal and fitness cost to variance in gene drive performance. Our findings reveal how spatial outcomes are affected by key parameters, and how this sensitivity varies tremendously between different gene drives. These spatial properties can be used to classify gene drive behavior and are useful to determine suitability for a particular application.

## 1 Introduction

Gene drives are genetic constructs that spread within target populations faster than would be expected under normal Mendelian inheritance, even if the construct reduces the fitness of drive-bearing organisms [2]. Gene drives are broadly classified by their intended function: population suppression or replacement. In this work, we focus on gene drives in the latter category. Population replacement gene drives carry a payload that modifies drive-bearing organisms to mitigate or remove harmful traits, such as the capacity to transmit disease in mosquitoes [10]. The gene drive is used to genetically “replace” a target population with one for which all or most of the individuals carry the payload. Gene drives have demonstrated potential in both modeling and laboratory experiments, and have been successfully engineered in the dengue mosquito *Aedes aegypti* [41], the agricultural pest *Drosophila suzukii* [75], the malaria mosquito *Anopheles gambiae* [30], and the invasive house mouse *Mus musculus* [29].

Gene drive potential is tempered by the possibility of the genetic cargo spreading to populations beyond the intended control region. Caution has been a guiding principle of technological development, with spatial confinement being a primary concern [22]. To reduce the risk of escape, gene drives can be designed so that self-sustained spread of the payload allele only occurs after exceeding a frequency, or threshold, in the target population. For these threshold-dependent gene drives, populations outside the control region tolerate a limited influx of drive-bearing organisms without population replacement occurring. So long as the payload frequency remains below a certain threshold, self-sustained spread does not occur. Threshold-dependent gene drives (henceforth referred to as threshold drives) are also reversible: releasing wild-type organisms can push the payload frequency to sub-threshold levels, leading to removal of the modified population. Examples of threshold gene drives include chromosomal translocations [11], engineered genetic incompatibility [47], tethered homing [21], Cleave-and-Rescue [55], and CRISPR-Cas9 toxin-antidote systems [14, 15].

The dynamics of threshold gene drives are characterized by an intrinsic tension between the local spread of a gene drive and the possibility of escape beyond the control region [8, 17, 23]. Indeed, if the control region comprises multiple target populations, gene drive spread is necessary to establish itself within each population. Gene drive success relies on the construct becoming established in the control region and spreading no further. These outcomes are not interchangeable, however: spatial confinement to the control region does not guarantee local establishment and vice versa. The relationship between these outcomes is complex as the spatial spread of a gene drive depends on factors unique to the target species and the gene drive itself [53].

The most important factors that determine the spatial outcomes of a gene drive are fitness cost and target organism dispersal (sometimes referred to as migration, immigration, or gene flow) [40, 50, 51]. However, the contribution of these parameters to gene drive performance differs tremendously. Buchman et al. (2018) found the migration rate between populations to be more influential to the equilibrium frequency of chromosomal translocations than fitness cost [11]. Reid et al. (2022) found that payload fitness cost was the strongest factor affecting the persistence of a homing drive [62], while the results of [51] suggest that homing drive spread between populations depends on generational migration. Altrock et al. (2011) found that increasing migration rates can cause extinction of an underdominant allele by disrupting a migration-selection equilibrium and increasing payload frequency [3]. The probability of fixation was primarily determined by the fitness of mutant homozygotes. Marshall and Akbari (2018) hypothesized that differences in dispersal habits and fitness costs between *Drosophila simulans* and *Aedes aegypti* produced the observed differences in spread of *Wolbachia* (a bacterial endosymbiont that behaves similarly to a gene drive) [46]. Clearly, the spatial dynamics of gene drives, and thus performance variance, overwhelmingly depend on fitness cost and target organism dispersal.

Parameter uncertainty confounds these issues and can add further variability to gene drive performance. Fitness cost and organism dispersal are notoriously difficult to measure accurately [42]. Fitness costs differ between laboratory and wild settings, and in the wild can change over time with environmental conditions or in response to selection [52]. Target organism dispersal (both natural and human-assisted) is similarly multifactorial and depends on season, population density, wind direction, and the movement of commerce, to give a few possible examples. Finally, some variance in performance will always be present due to stochasticity. Our goal in the present work is to use techniques from global sensitivity analysis to measure how the performance of threshold gene drives depends on fitness cost and target organism dispersal. These sensitivity indices are used alongside other metrics to investigate the risk of failure for different threshold drives.

Modeling studies that investigate the confinement properties of threshold drives often use deterministic models or make simplifying assumptions such as unidirectional or fixed dispersal, or infinite population size [3, 4, 11, 13, 14, 17, 21, 24, 35, 43]. Such studies rarely account for parameter uncertainty in dispersal and payload fitness cost and their interacting influence on threshold drive performance. Stochasticity is similarly neglected. We briefly discuss several past studies investigating the confinement and permanence properties of threshold (or otherwise spatially-limited) gene drives. Marshall and Hay (2012) used metapopulation models to compare the confinement properties of different replacement drives [45]. Their analysis includes a two-population model with bidirectional dispersal between control and non-control regions. A stochastic version of this model is used to compute the probability of invasion of the non-control region over a period of time. Similar to Marshall and Hay, Dhole et al. (2017) compared the localization and permanence of spatially self-limiting gene drives [23]. They use a two-patch model and assume a large population with non-overlapping generations. For the simulations with bidirectional dispersal, they calculate the mean payload frequency in both patches as migration rate and introduction frequency vary. This analysis is repeated for different payload fitness costs. Finally, Sanchez et al. (2020) use a spatially-explicit model to investigate confinement and remediation of two threshold gene drives in an Australian suburb [66]. The authors conduct a sensitivity analysis that perturbs the fitness cost and dispersal parameters by fixed amounts. We extend this body of work with a discrete-time mathematical model that incorporates space, stochasticity, and parameter uncertainty. We use computer simulations to understand how spatial outcomes depend on key model parameters and conduct a variance-based sensitivity analysis to compare the influence of fitness cost and organism dispersal to changes in gene drive performance. We pay particular attention to gene drive failure and compute failure probabilities for each threshold drive within their respective parameter ranges.

In our model, we assume the target population is comprised of demes, or patches. Mating is panmictic within each patch, and organism dispersal is bidirectional between adjacent patches. We consider two different forms of dispersal: short- and long-distance. Short-distance dispersal occurs between adjacent patches within the control region, while long-distance dispersal occurs between control and non-control regions. To parameterize our system, we take the life history and dispersal parameters to be that of the dengue mosquito, *Aedes aegypti*. *Ae. aegypti* transmits viruses responsible for causing illness in over 100 million people annually [67], including Zika, Chikungunya, West Nile, and dengue. We conduct our analysis over a wide range of parameter values to reflect uncertainty in parameter estimates [74]. We investigate the spatial outcomes of four threshold gene drives: two-locus underdominance [19], tethered homing [21], toxin-antidote recessive embryo (TARE) [15], and toxin-antidote dominant embryo (TADE) [14].

The objectives in this manuscript are twofold: (i) to understand how payload fitness cost and organism dispersal affect spatial outcomes, including risk of failure, and (ii) to quantify sensitivity of gene drive performance to these parameters. Regarding (i), it is presently unclear the extent that the spatial performance of threshold drives relies on target organism dispersal and fitness cost, at least within the same modeling framework that allows for meaningful comparison between threshold drives. By investigating how either parameter contributes to drive failure, these findings can improve monitoring efficiency. Our second objective, (ii) to quantify sensitivity of threshold drive performance to either parameter, is critical in designing effective risk management strategies, choosing the most suitable threshold drive for a specific application, and capturing the full scope of possibilities associated with transgenic release [8, 50, 63, 74].

## 2 Gene drives

We analyze the performance of four threshold gene drives: two-locus underdominance [19], tethered homing [21], toxin-antidote recessive embryo (TARE) [15], and toxin-antidote dominant embryo (TADE) [14]. We choose these gene drives for their diversity in mechanism and dynamics. A list of genotypes for each gene drive and their fitnesses relative to wild-type is provided in Table SM1.

### 2.1 Two-locus underdominance

Underdominance refers to when heterozygotes are less fit than either parental homozygote [33]. Confinement occurs because drive homozygotes that disperse into predominantly wild-type populations will give rise to progeny with low fitness. Engineered two-locus underdominance typically relies on lethal-suppressor mechanisms [1, 19, 43]. (Here, we intentionally avoid use of the phrase “toxin-antidote” so as to avoid confusion with the toxin-antidote gene drives considered later in this paper.) In engineered underdominance, each transgenic allele possesses a lethal gene that is suppressed by the complementary allele. Let *AABB* be the genotype of wild-type individuals. We assume that the underdominance transgenes are unlinked and inserted at the *A* and *B* loci [19]. Let α and β denote the underdominance transgenes, so that the suppressor gene in the α allele rescues the organism from the lethal β gene and vice versa. We assume that transgenes are strongly suppressed, or that a transgenic individual must carry at least one of each transgene to be viable [19]. Viable organism genotypes are *AABB* (wild-type), *A*α*B*β, *A*αββ, αα*B*β, and ααββ. A feature of two-locus underdominance is that it possesses an intrinsic threshold (approximately 27% [25]), even in the absence of a fitness cost [19, 43].

### 2.2 Tethered homing

Homing gene drives rely on Cas9-gRNA complexes to cut the homologous chromosome and exploit DNA repair mechanisms to duplicate payload alleles [26, 27]. This process converts heterozygotes to homozygotes. Homing drives can spread from small releases, in contrast to threshold drives which must be present above a certain frequency to propagate. This is a problem for spatial confinement [51]. Dhole et al. (2018) proposed a “tethered drive” that couples a homing payload with a threshold drive; the latter acting as a “tether” to limit payload spread [21]. Homing activity is conditional on a drive-bearing organism possessing both the payload and the tether, so that self-propagating spread of the payload only occurs in those organisms that carry the threshold drive components.

As in Dhole et al. (2018), we take the tether of our threshold drive to be two-locus underdominance. Let *AABBCC* be the genotype of wild-type individuals. The underdominance drive is inserted at the *A* and *B* loci, and the homing drive is inserted at the *C* locus. We maintain the previous assumption that the transgenes are strongly suppressed so that viable transgenic organisms must possess at least one underdominance component at the *A* and *B* loci. The viable genotypes are the same as those provided for two-locus underdominance in the previous section, regardless of the *C* locus: *AABB*, *A*α*B*β, *A*αββ, αα*B*β, and ααββ.

### 2.3 Toxin-antidote systems: TARE and TADE

We consider both toxin-antidote recessive embryo (TARE) and toxin-antidote dominant embryo (TADE). Both systems rely on a Cas9-gRNA complex to disrupt a target site (the “toxin”) [15]. In the case of TARE, the broken allele is recessive lethal so the target site is haplosufficient (i.e., viable organisms have at least one unbroken allele). For TADE, the broken allele is dominant lethal so the target site is haploinsufficient (i.e., viable organisms have two unbroken alleles). The payload allele in both cases includes a recoded copy of the target allele that is unrecognizable by the Cas9-gRNA complex (the “antidote”). Drive-bearing individuals can still be viable because of the recoded target allele. TARE and TADE exhibit threshold-dependent dynamics when the payload allele imposes a sufficiently high fitness cost [14, 15]. Let α denote the transgene, *A* the wild-type allele, and β the broken allele. Genotype ββ is the only non-viable genotype for TARE. For TADE, the non-viable genotypes are αβ, *A*β, and ββ.

## 3 Model

We use a discrete-time patch model with sex and age structure to investigate how spatial outcomes of threshold drives depend on fitness cost and organism dispersal. Each patch represents a panmictic population. We use a multi-patch model, as it is common for a species to form geographically distinct “colonies” owing to environmental heterogeneities and organism characteristics [22, 37, 68]. For example, houses act as units of clustering for the mosquito *Ae. aegypti* [28], and herbivorous pests cluster around crop monoculture. In this section, we describe the dynamics within and between patches, the model used for simulation, and the stochastic version of the model.

### 3.1 Within-patch dynamics

Populations are diploid and assumed to be well-mixed within a patch. There are four compartments in total, distinguished by age and sex: juvenile males, juvenile females, adult males, and adult females. We use a hat (^) to refer to juvenile male parameters and variables when appropriate. Let *Ĵ_G_*(*i*, *t*) denote the number of male juveniles with genotype *G* in patch *i* at time *t*. Similarly, let *J_G_*(*i*, *t*) denote the number of female juveniles with genotype *G* in patch *i* at time *t*. For simplicity, we let *J* (without a subscript) denote the total juvenile population at a specific time and patch, i.e.

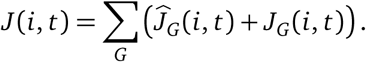

Juvenile survival depends on juvenile density and is denoted by the function *f* [*J*(*i*, *t*)]. This is

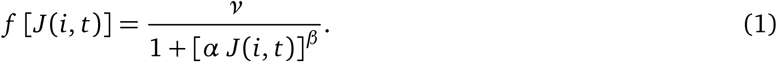

The probability of density-independent survival is given by *v*. The parameter β determines the strength of density dependence exhibited by the target organism population, while α adjusts the population size at equilibrium. We set β = 2, which corresponds to strong density dependence (as exhibited by *Ae. aegypti* [12]), and α = 1.7 × 10^−4^ for an equilibrium population of 2000 individuals. We choose Eqtn. (1) as our functional form of density-dependent mortality for its flexibility in capturing density-dependent processes [9]. Juveniles emerge as adults with probability η per time step. Juvenile hatching and emergence depends on their genotype and sex. The hatching probability is multiplied by an early-acting relative fitness term, denoted by ϕ̂*_G_* and ϕ*_G_* for males and females, respectively. Additionally, the emergence probability is multiplied by a late-acting relative fitness term, denoted by ψ̂*_G_* and ψ*_G_* for males and females, respectively. The values of these parameters depend on when the fitness cost of the gene drive is imposed. For wild-type organisms, we take these fitness parameters to be unity. Gene drive fitness costs are discussed in greater detail below.

Let *M_G_*(*i*, *t*) denote the number of male adults with genotype *G* in patch *i* at time *t*, and similarly *F_G_*(*i*, *t*) denotes the number of female adults with genotype *G* in patch *i* at time *t*. Adult females lay λ eggs per day. Adult mortality depends on sex, with male and female adults dying with probabilities µ*_M_* and µ*_F_*, respectively, per time step. Within-patch dynamics for each population are captured by the following system of equations:

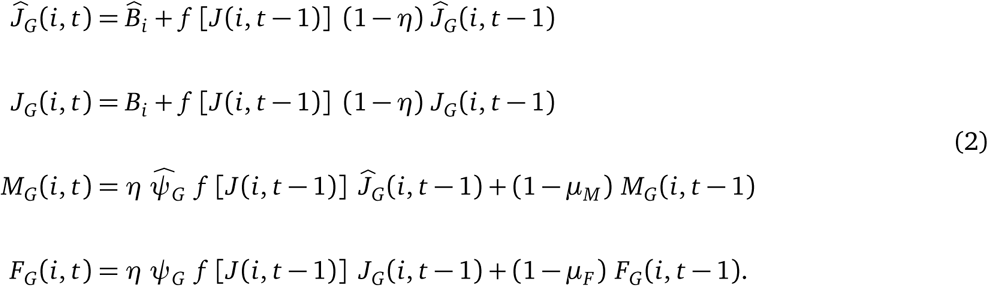

Birth terms in patch *i* for male and female juveniles are given by *B̂_i_* and *B_i_*, respectively. These are

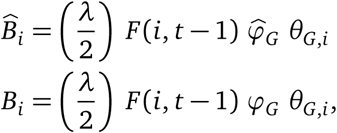

where θ*_G_*_,*i*_ is the probability that a genotype *G* offspring results from parents mating in patch *i*. To compute θ*_G_*_,*i*_, we sum over all possible parental genotype combinations weighted by their frequencies. To each such combination of parental genotypes corresponds a conditional probability, *P*(*G*|*H*, *K*), or the probability that a genotype *G* offspring is produced from a genotype *H* mother and a genotype *K* father. The probability of mating with a genotype *K* male is just the proportion of genotype *K* in the male population, as mating is random. We assume there is no difference in mating ability between genotypes. Taken together, this gives

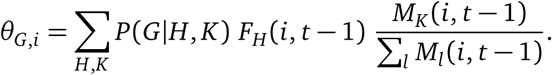

Transgenic organisms may incur a fitness cost due to each of the gene drive alleles, such as might be caused by leaky toxin expression or gene interactions with the modified locus. All fitness costs are taken to be multiplicative and affect both sexes equally. Fitness costs are early- or late-acting depending on whether they occur before or after density-dependent mortality, respectively. In Eqtn. (2), the early-acting relative fitness of an organism is the probability that it hatches, while the late-acting relative fitness is the probability it ecloses as an adult following pupation. We denote the fitness cost of the gene drive by *c* so that individuals with *n* transgenes have relative fitness (1 − *c*)*^n^*. Parameter definitions, values, and their corresponding references are provided in Table 1.

**Table 1:** Parameter definitions and values for the within- and multi-patch models (Eqtns. (2) and (3), respectively) of gene drive spread in a population of *Ae. aegypti* mosquitoes.

| Parameter | Definition | Value | Reference |
| --- | --- | --- | --- |
| $\lambda$ | Daily egg-laying rate per adult female | 15.06 | [18] |
| $\hat{\varphi}_G (\varphi_G)$ | Early-acting relative fitness of males (females) with genotype $G$ | varies | N/A |
| $\hat{\psi}_G (\psi_G)$ | Late-acting relative fitness of males (females) with genotype $G$ | varies | N/A |
| $\eta$ | Daily juvenile emergence probability | 0.1 | [18] |
| $\nu$ | Daily density-independent juvenile survival probability | 0.9351 | [76] |
| $\alpha, \beta$ | Density-dependent competition parameters | $1.7 \times 10^{-4}, 2$ | [69] |
| $\mu_M (\mu_F)$ | Daily survival probability for male (female) adults | 0.6930 (0.8249) | [71] |
| $m_{ij}$ | Daily short- (long-) distance dispersal probability from patch $i$ to patch $j$ | 0.1305 ($4.0 \times 10^{-4}$) | [31, 69] |
| $c$ | Fitness cost of each transgene | varies | N/A |

### 3.2 Multi-patch model

The multi-patch system differs from that in Eqtn. (2) in that it includes movement between patches. Dispersal probabilities are given by a matrix with entries *m_i j_* denoting the daily dispersal probability from patch *i* to *j*. For a system with *P* patches, this is

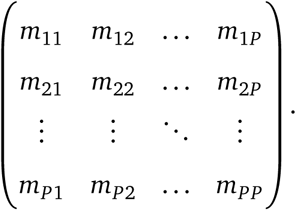

The probability that an adult remains in patch *i* is *m_ii_* and, because we assume that no individuals are lost from dispersal, we have that 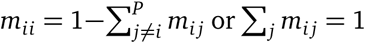. We assume that *m_i j_* = *m_ji_* for all (*j*, *i*) ∈ *P* × *P*. At equilibrium, the populations within each patch remain constant on average. We assume that only adults disperse. The sequence of model events for each time step is as follows: mating, adult dispersal, oviposition, and mortality. This new system is

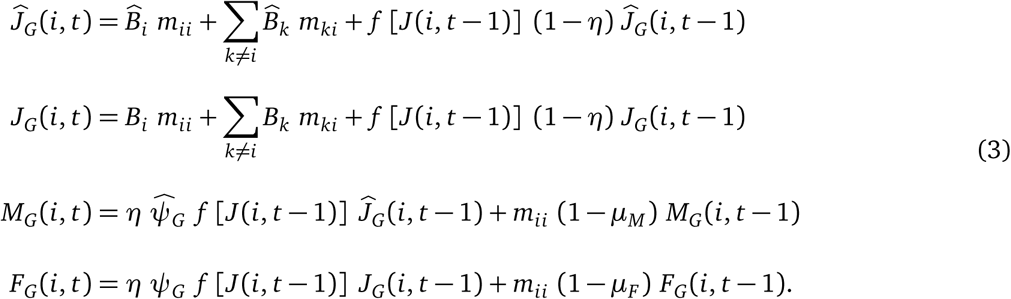

The birth terms in Eqtn. (3) now account for dispersal, as adult females can mate and oviposit in different patches.

### 3.3 Three-patch system

We use a multi-patch model to investigate the ability of a gene drive to spread within and remain confined to a control region. We wish to explore the general relationship between gene drive spread and confinement, as determined by key model parameters, and not the dynamics associated with a specific location. In a hypothetical release, there will be a barrier inhibiting target organism dispersal between the control and non-control regions [50]. In practice, such barriers include bodies of water or broad stretches of land, but anthropological barriers (e.g., highways) have also been shown to inhibit insect dispersal [34, 73]. The barrier inhibits but does not prevent dispersal [8, 42, 50, 72]. Furthermore, we are interested in simulating spread over short distances within the control region.

These considerations lead us to the three-patch system depicted in Fig. 1. This system is comprised of three populations assorted in a linear array with bidirectional dispersal between adjacent populations. Patches 1 and 2 form the control region, while patch 3 represents the non-control region. We refer to dispersal between patches 1 and 2 as **short-distance dispersal**, and refer to the exchange of organisms between patches 2 and 3 as **long-distance dispersal**. Short-distance dispersal occurs more frequently than long-distance dispersal due to the barrier between the control and non-control regions. We assume short-distance dispersal is at the scale of 100-200 m., or an agreeable distance between villages or other human habitations at which *Ae. aegypti* populations would cluster [49]. Daily dispersal estimates for *Ae. aegypti* at this scale suggest the daily probabilities fluctuate between 1 − 20% [69]. Long-distance dispersal probabilities are computed assuming a barrier of at least 500 m. in width, representing the upper limit of *Ae. aegypti* dispersal and thus what might be considered practically suitable for a wild release [31].

**Figure 1:**
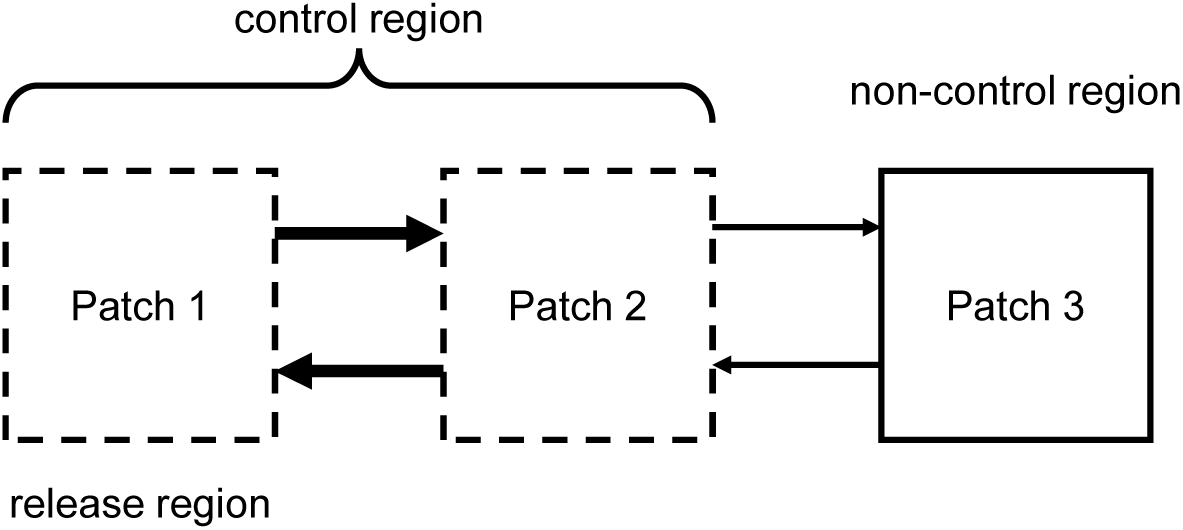
Schematic of the three-patch model. Patches 1 and 2 comprise the control region (broken boxes) while patch 3 represents the non-control region (solid box). Gene drive release occurs in patch 1. Arrows indicate dispersal rate and direction, while arrow thickness denotes frequency (thicker arrows indicate higher frequency). Dispersal is bidirectional and either short-distance (between patches 1 and 2) or long-distance (between patches 2 and 3).

The ideal outcome following transgenic release in patch 1 is that the payload reaches a high and persistent frequency in patches 1 and 2, but does not spread to patch 3. However, this represents only one of four possible outcomes following transgenic release:

1. The gene drive fails to persist in patch 1 and goes extinct.
2. The gene drive persists in patch 1 but does not spread to patch 2.
3. The gene drive establishes in patches 1 and 2 but does not spread to patch 3.
4. The gene drive spreads to all patches.

Scenario 3 is the ideal outcome, while the remaining cases represent different forms of gene drive failure. Scenarios 1 and 2 have received little attention relative to Scenario 4. Our analysis suggests that target organism dispersal over short distances is typically high enough to prevent Scenario 2, so we ignore it. This leaves Scenarios 1 and 4 as the only forms of drive failure we consider. For brevity, we refer to Scenario 1 as **extinction** (failure in establishment) and Scenario 4 as **escape** (failure in confinement).

### 3.4 Stochastic model

We use a stochastic version of the three-patch model to capture the effects of random events. Male-female pairings are determined by a multinomial distribution with male mates apportioned based on their frequency in the population at that time. Dispersal, progeny sex, and death are binomially distributed. Oviposition occurs throughout a female’s life and begins as soon as she mates. The daily number of progeny produced per adult female is Poisson distributed with mean λ. Offspring genotypes are multinomially distributed and based on zygote frequencies calculated from parental genotypes. Both juvenile survival and emergence are binomially distributed.

## 4 Methods

### 4.1 Stochastic simulations

We perform Monte Carlo simulations to investigate how the payload frequency of the gene drive changes with stochastic variation and for different parameter values. These results are used to compute the failure probability for each gene drive, and the mean and variance of the drive allele frequency at certain time points post-release. The variance is a measure of the frequency of dichotomous outcomes (extinction and fixation) and is used to compute sensitivity indices of drive frequency with respect to key parameters.

Short- and long-distance dispersal rates are assumed to be uniformly distributed over the ranges [0.01, 0.2] and [10^−6^, 10^−3^], respectively. (We consider non-uniformly distributed dispersal parameters in the Appendix.) These ranges were chosen to include the nominal values calculated for *Ae. aegypti* while accommodating natural variation or uncertainty. Fitness cost values are chosen based on simulated performance of the gene drive under nominal dispersal conditions. For each gene drive, the range of fitness cost values used for the Monte Carlo simulations are those for which the probability of gene drive failure did not exceed 10%. We refer to this range of fitness cost values as the **fitness profile** of the gene drive. This choice is conservative, as laboratory-measured fitness costs are uncertain, and fitness effects will naturally differ in the wild than those measured in the laboratory [8]. Monte Carlo sampling is performed via a Latin hypercube routine [58]. A positive payload frequency is expected in the non-control region due to long-distance dispersal, so we take escape to be when this frequency exceeds 10%.

Gene drive release occurs after 100 days to allow the wild-type populations in the system to stabilize. Gene drives are introduced with four weekly 6:1 releases. We use large releases to ensure that the threshold drives successfully invade over a wide range of parameter space. The large releases also help to avoid the prolonged invasion period that can occur for low introduction frequencies, or introduction frequencies near a bistable equilibrium. The genotype of the introduced transgenic organisms depends on the gene drive. We use Greek letters to indicate transgenes and Roman letters to indicate wild-type alleles. Drive homozygotes are released for two-locus underdominance (genotype ααββ), while heterozygotes are released for the toxin-antidote drives (genotype *A*α). For the tethered homing gene drive, we use the two-genotype release described in [21]. Specifically, 95% of individuals in the first release are homozygous for the underdominance components and lack the homing component (ααββ *CC*) while the remaining 5% of individuals are heterozygous for the homing component (ααββ *C*γ). In the remaining three releases, all transgenic organisms are homozygous for the underdominance component and lack the homing component (ααββ *CC*). For tethered homing, we fix the fitness cost of the underdominance transgene at 0.05 and vary only the fitness cost of the homing payload allele. The tethered homing gene drive includes a homing component that relies on cleaving the homologous chromosome and repairing the break via homology-directed processes. We assume that the probability of homology-directed repair is 100%. (For *Ae. aegypti*, this is not far from what has been achieved experimentally [5, 41].)

### 4.2 Variance-based sensitivity analysis

We conduct a variance-based sensitivity analysis to investigate how gene drive performance is sensitive to changes in payload fitness cost and target organism dispersal. For each gene drive, we compare the relative influence of dispersal and fitness cost on spatial outcomes, and quantify the contribution of each parameter to performance variance, or uncertainty. We measure gene drive performance by the transgene frequency at yearly intervals after release.

Let 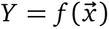 denote a quantity of interest for some model *f* and input vector 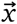. The first-order sensitivity index of parameter *x_u_* is [64]

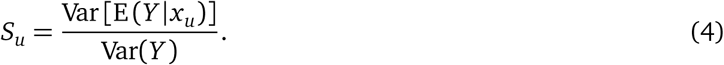

First-order indices identify the parameter that induces the largest reduction in outcome variance Var(*Y*) if its value was known accurately [7]. That is, the first-order indices tell us which individual parameters, if fixed, reduce uncertainty in *Y* most substantially. For example, *S_u_* = 0.5 means that fixing parameter *u* reduces Var(*Y*) by 50%. Note that the sensitivity indices given by Eqtn. (4) are relative: fixing parameters to reduce Var(*Y*) is pointless if Var(*Y*) ≈ 0.

The total-order sensitivity index accounts for all contributions of the parameter *x_u_* to outcome uncertainty, including higher-order interactions between other parameters [65]. The total-order sensitivity index of *x_u_* is [64]

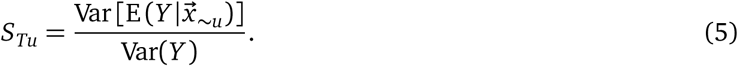

Here, 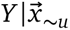 means conditional on every parameter in 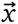 except for *x_u_*, so that all parameters except for *x_u_*are fixed. We adopt the convention of others (e.g., [77]) that parameter *x_u_* has a significant effect on *Y* if *S_Tu_* > 0.05. The difference between the total- and first-order indices, *S_Tu_* −*S_u_*, measures the extent of higher-order interactions [64]. A large difference indicates that most of the contribution of *x_u_* to the variance of *Y* depends on the values of other parameters.

We use the estimators proposed in [7] to numerically compute the indices in Eqtns. (4) and (5). Let ***A*** and ***B*** denote sample matrices with dimension *N* × *k*, where *N* is the number of Monte Carlo simulations and *k* is the number of model parameters. Each matrix row is a vector of parameter samples from their respective distributions. We form the matrix ***A****_u_* by replacing column *u* in ***A*** with the corresponding column in ***B***, and similarly for ***B****_u_*. Let the vector 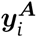 denote the output of a stochastic model evaluation using the parameters in the *i*th row of ***A***. The first-order indices of *x_u_* are

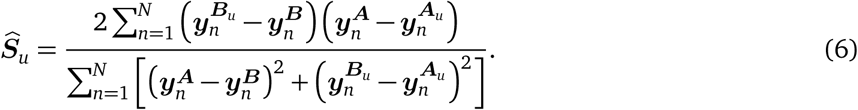

The total-order indices of *x_u_* are estimated as

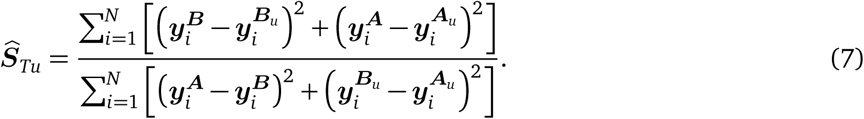

In the context of our problem, the quantity of interest is the gene drive allele frequency in patch *i* at a certain time point post-release. Unless otherwise mentioned, we chose four years post-release as this was sufficient time for payload frequencies to stabilize in each patch. Our model has *k* = 3 parameters that we explore in this analysis: the payload fitness cost, the short-distance dispersal probability, and the long-distance dispersal probability. We choose *N* large enough to achieve convergence of the estimators given in (6) and (7). This is validated graphically by the plots shown in Figs. SM3-SM6.

We use bootstrapping to compute confidence intervals for the numerical estimates of the first- and total-order indices [54]. To do this, we sample from the *N* model simulations with replacement to recalculate the sensitivity indices. This process is repeated 1000 times until a distribution is obtained, from which confidence intervals are computed. The 95% confidence intervals computed for first- and total-order indices are provided in Tables SM4 and SM5, respectively.

## 5 Results

It was always the case that fitness cost and short-distance dispersal significantly affected the allele frequency in the control region (i.e., *S_Tu_* > 0.05). Similarly, it was always the case that fitness cost and long-distance dispersal significantly affected payload frequency in the non-control region. We reserve comment for cases that deviate from these patterns (e.g., short-distance dispersal significantly affecting the frequency in the non-control region). We report both the variance (σ̂^2^) and standard deviation (σ̂) of the payload frequency in the control and non-control regions. For the control region, we report a single variance obtained by taking the mean of the allele variance for patches 1 and 2.

### 5.1 Two-locus underdominance

Two-locus underdominance is highly efficient at carrying a deleterious payload into a population. Fig. 2 shows the failure probability of an underdominance drive as fitness cost varies. The fitness profile of underdominance, or the range of fitness cost values for which the failure probability did not exceed 10%, is *c* ∈ [0, 0.22]. Introgression is successful despite the high costs of the genetic cargo: for example, when *c* = 0.20, drive-heterozygotes (genotype *A*α*B*β) have relative fitness (1 − *c*)^2^ = 0.64. Beyond *c* = 0.21, the extinction probability rapidly increases to 100% due to the fitness cost imposed on heterozygotes becoming unsustainable for drive introgression. In the parameter regime near *c* = 0.21, slight changes to the payload fitness cost can dramatically increase the probability of failure [8]. Two-locus underdominance also remains confined to the control region, as we never observed escape across the conditions explored in the Monte Carlo simulations. Confinement occurred even for simulations with no payload fitness cost (*c* = 0).

**Figure 2:**
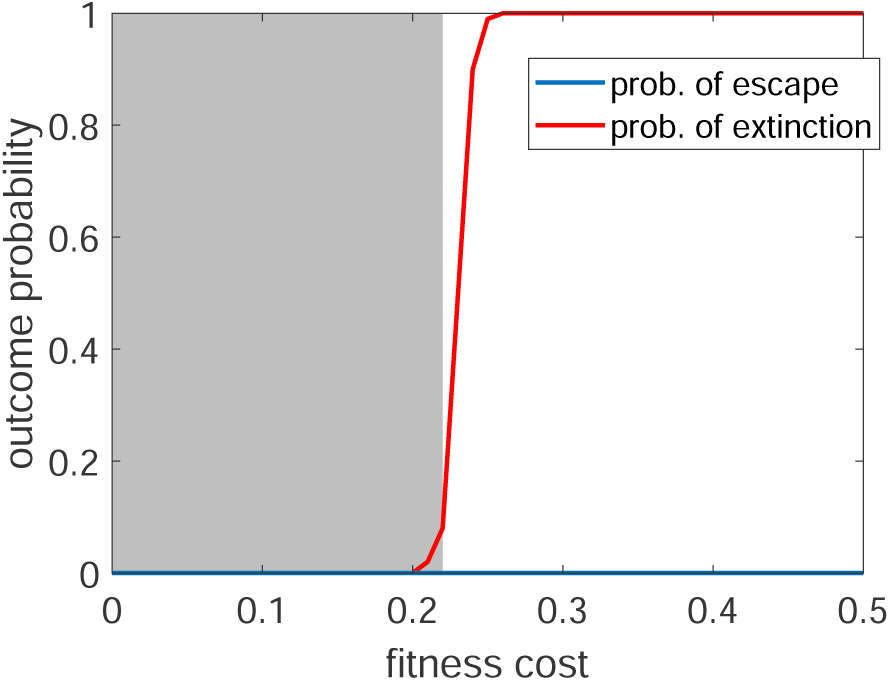
The failure probability for a two-locus underdominance drive as fitness cost varies. Failure is classified as either failure in persistence (extinction; red line) or failure in confinement (escape; blue line). The shaded region denotes the fitness profile of the gene drive, or the range of fitness cost values for which the failure probability is below 10%, which is *c* ∈ [0.00, 0.22]. Note that the probability of escape lies along the *x* −axis as it was never the case that the gene drive escaped the control region in our simulations. Dispersal parameters are fixed at their nominal values: the daily probability of short- and long-distance dispersal is approximately 13% and 0.04%, respectively (see Table 1). Each data point is the average of 100 stochastic simulations.

Payload frequency in the control region varied substantially for two-locus underdominance. This indicates that the payload frequency is sensitive to our choice of dispersal and fitness cost parameters. The standard deviation of the allele frequency at equilibrium was σ̂ = 28.5% (σ̂^2^ = 8.1%; see Figs. 3(a)-(b)). Increasing the fitness cost of an underdominance gene drive reduces the payload frequency at equilibrium. However, the first-order sensitivity indices reveal that the payload frequency in the control region is slightly more sensitive to changes in short-distance dispersal than fitness cost (32.7% versus 25.6%; see Table 2). This sensitivity of performance to dispersal has been found by other investigators. Leftwich et al. (2018) found that dispersal rates can cause underdominant mutations to struggle in spreading through metapopulations [39]. Altrock et al. (2011) studied an underdominant mutant allele in a two-population model and found that high migration increases allele frequency variance, and that allele dynamics (including extinction) strongly depend on the migration rate [3]. Managing organism dispersal, especially over short distances, is the most effective strategy to control payload variance of a two-locus underdominance gene drive.

**Figure 3:**
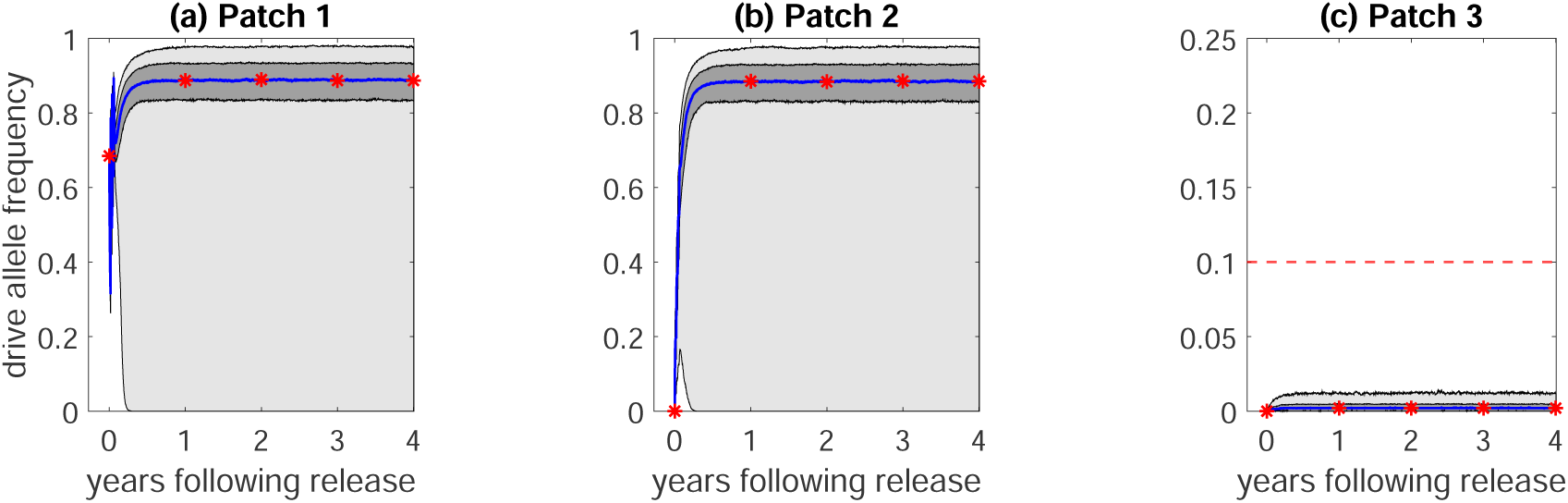
Stochastic simulations of a two-locus underdominance gene drive in the three-patch system. The middle 50th and 95th percentiles of the payload frequency are given by the light and dark gray regions, respectively. The median trajectory is denoted by the blue line with red asterisks. The plots show payload frequencies in **(a)** patch 1, **(b)** patch 2, and **(c)** patch 3. These computations are based on 1000 stochastic simulations. Note the change in *y*-axis in (c); gene drive escape occurs if the payload frequency exceeds 10%, denoted by the broken red line.

**Table 2:** The first-order indices, computed using Eqtn. (6), for each parameter in the case of two-locus underdominance (TLU), two-locus underdominance tethered homing (TLUTH), toxin-antidote recessive embryo (TARE), and toxin-antidote dominant embryo (TADE). The indices for TARE were computed using the drive allele frequency six years post-release; all other drive indices are computed using the payload frequency four years post-release. 95% confidence intervals for these indices are provided in Table SM4.

| Gene drive | Parameter | Patch 1 (%) | Patch 2 (%) | Patch 3 (%) |
| --- | --- | --- | --- | --- |
| TLU | Fitness cost | 26.49 | 24.57 | 37.64 |
|  | Short-distance dispersal | 29.96 | 34.62 | 3.05 |
|  | Long-distance dispersal | 0.08 | 0.12 | 41.58 |
| TLUTH | Fitness cost | 0.57 | 0.20 | 57.91 |
|  | Short-distance dispersal | 76.19 | 90.62 | 0.93 |
|  | Long-distance dispersal | 0.02 | 0.11 | 21.78 |
| TARE | Fitness cost | 30.57 | 30.46 | 17.29 |
|  | Short-distance dispersal | 5.72 | 6.32 | 0.00 |
|  | Long-distance dispersal | 0.13 | 0.23 | 9.47 |
| TADE | Fitness cost | 11.63 | 10.24 | 15.66 |
|  | Short-distance dispersal | 0.71 | 4.38 | 0.00 |
|  | Long-distance dispersal | 0.07 | 0.24 | 5.60 |

The high contribution of short-distance dispersal to payload variance also means that unfavorable dispersal conditions are more likely than high payload costs to lead to failure. This is supported by our Monte Carlo analysis: the gene drive went extinct in 11% of simulations, all of which were characterized by low short-distance dispersal rates (less than 10% probability per day) and higher fitness costs (*c* ≥ 0.08; see Fig. SM1(a)). Studies in continuous space have similarly shown that it can be difficult for underdominance systems to successfully persist in the control region [17]. Altrock et al. (2011) found that extinction was the most common outcome for an underdominant allele under different fitness and dispersal scenarios following system disturbance at an interior equilibrium [3].

Confinement of the underdominance drive is robust to changes in fitness cost and dispersal (see Fig. 3(c)). Escape did not occur, even for high rates of long-distance dispersal and no fitness cost (*c* = 0). Other investigators have found that establishment in neighboring populations can occur for high dispersal probabilities per generation, depending on population size [45]. The first-order indices suggest that payload variance in the non-control region is more sensitive to changes in long-distance dispersal than fitness cost (41.6% compared to 37.7%). As with payload persistence in the control region, managing target organism dispersal is key to achieving gene drive confinement. Interestingly, short-distance dispersal significantly affected the payload frequency in the non-control region (see Table SM2). Thus, dispersal over short distances is important for the spatial dynamics of two-locus underdominance, even at larger scales.

Up until this point, we have measured sensitivity of payload frequency to model inputs assuming only a single parameter changes at a time. Higher-order interactions quantify the effect of modifying multiple parameters simultaneously on gene drive performance, obtained by taking the difference between total- and first-order indices, *Ŝ_Tu_* − *Ŝ_u_*. These are given in Table SM2. While a majority of payload variance is due to changes in individual parameters, higher-order interactions are still present. We have already seen why: higher short-distance dispersal is required to ensure adequate drive introgression for high payload fitness costs. For low or moderate fitness costs, manipulating dispersal alone is sufficient to ensure ideal spatial outcomes.

### 5.2 Tethered homing

A tethered homing drive can carry a payload to high frequency within the control region across a large range of fitness costs [21]; the fitness profile of tethered homing is *c* ∈ [0.04, 0.38] (see Fig. 4).

**Figure 4:**
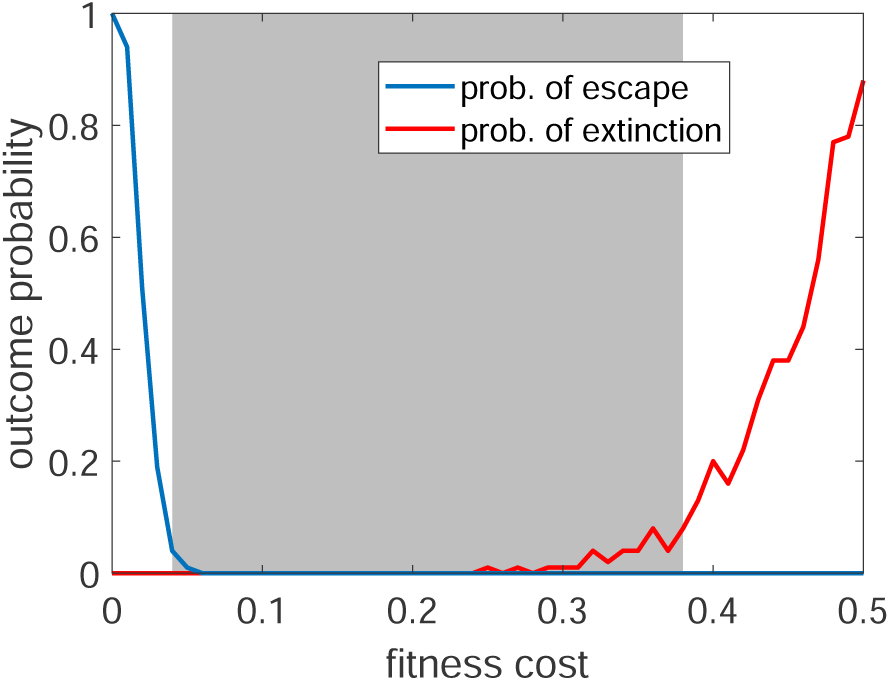
The failure probability for a tethered homing drive as fitness cost varies. Failure is classified as either failure in persistence (extinction; red line) or failure in confinement (escape; blue line). The shaded region denotes the fitness profile of the gene drive, or the range of fitness cost values for which the failure probability does not exceed 10%, which is *c* ∈ [0.04, 0.38]. Fitness cost refers to the cost of the homing payload and not of the underdominance components, which are fixed at 0.05. Dispersal parameters are fixed at their nominal values: the daily probability of short- and long-distance dispersal is approximately 13% and 0.04%, respectively (see Table 1). Each data point is the average of 100 stochastic simulations.

This trait is shared with the underdominance drive considered above. However, unlike underdominance, tethered homing requires a minimum fitness cost to prevent gene drive escape. When *c* < 0.04, selection against the payload in the non-control region is low enough that it accumulates over time. Spread of the payload allele in the non-control region is facilitated by Mendelian inheritance and not by gene drive, as the underdominance components of the tethered homing drive remain confined to the control region. This is an interesting example of a gene drive component (in this case, the homing payload) building up in a non-control region in the absence of drive-induced super-Mendelian inheritance.

As payload fitness costs increase, the extinction probability climbs gradually rather than suddenly (compare red curves in Figs. 4 and 2). This is because payload spread in a population for which most individuals carry the underdominance component behaves like a standard homing drive. If the payload fitness cost is sufficiently high, introgression depends on the initial frequency of the gene drive [20]. Consequently, the probability of extinction increases gradually with fitness cost. Payload extinction is remedied by increasing the frequency of payload-bearing organisms (genotype ααββ *C*γ) [21].

Payload variance in the control region was moderate for the tethered homing gene drive: the standard deviation of the payload frequency in the Monte Carlo simulations was σ̂ = 14.7% (σ̂ = 2.2%; see Fig. 5(a)-(b)). This variation is primarily due to extinction events, which occurred in 2.7% of simulations. Extinction in this case refers to disappearance of the homing payload component and does not necessarily refer to disappearance of the underdominance component. Drive extinction occurred more frequently for simulations with low short-distance dispersal rates, typically less than 6% per day. In contrast, extinction occurred over a wide range of payload fitness costs (see Fig. SM1(b)). These observations are confirmed by the first-order indices, which indicate that the performance of the tethered homing drive in the control region is overwhelmingly determined by short-distance dispersal, accounting for 83.4% of payload variance (see Table 2). Long-distance dispersal and fitness cost each accounted for less than 1% of payload variance. The tethered drive achieved near-fixation in the control region for most simulations, even for high payload fitness costs (see Figs. 5(a)-(b)). This is desirable when few unmodified organisms can still inflict serious injury, such as disease-competent vectors in a fully susceptible human population [61].

**Figure 5:**
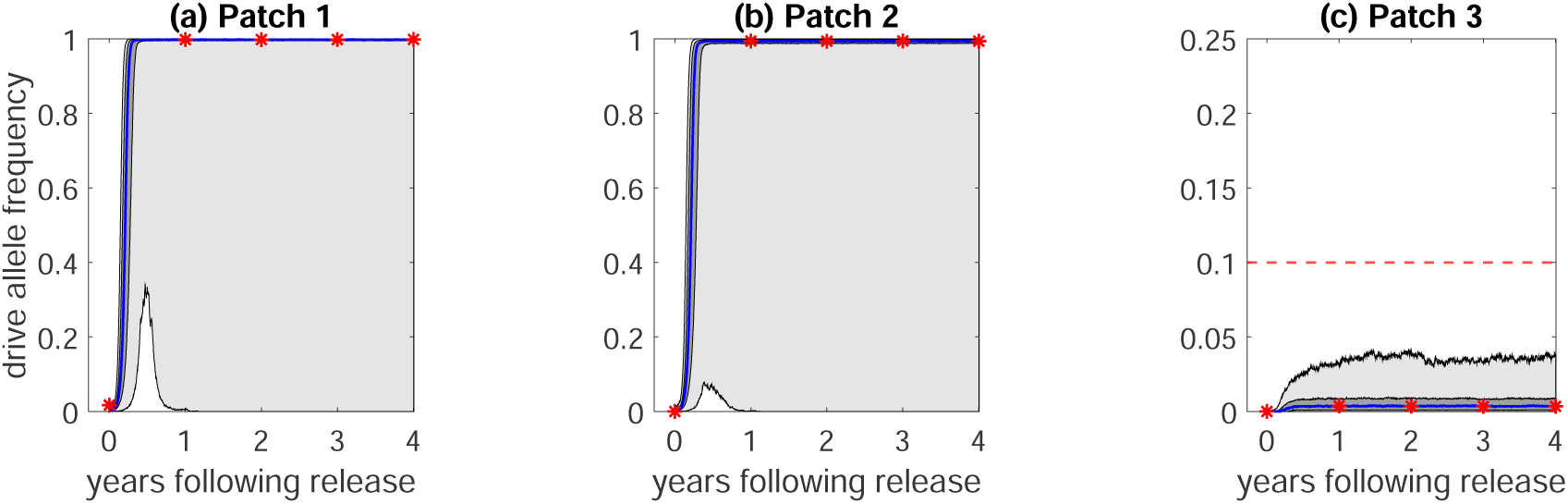
Stochastic simulations of a tethered homing gene drive in the three-patch system. Only the frequency of the (homing) payload allele is shown. The middle 50th and 95th percentiles of the payload frequency are given by the light and dark gray regions, respectively. The median trajectory is denoted by the blue line with red asterisks. The plots show payload frequencies in **(a)** patch 1, **(b)** patch 2, and **(c)** patch 3. The fitness cost of the underdominance transgene is 0.05, while the fitness cost of the homing component varies. These computations are based on 1000 stochastic simulations. Note the change in *y*-axis in (c); gene drive escape occurs if the payload frequency exceeds 10%, denoted by the broken red line.

Spatial confinement of the payload was insensitive to changes in dispersal and fitness cost parameters, so that payload variance in the non-control region was small. The computed standard deviation was σ̂ = 1% (σ̂^2^ = 0.01%; see Fig. 5(c)). Escape was rare and occurred in only 0.8% of simulations. Simulations producing escape had low payload fitness costs (*c* < 0.05) and long-distance dispersal probabilities exceeding 0.05% (see Fig. SM2). Escape occurs because payload alleles accumulate in the non-control region and not because of super-Mendelian propagation. Dhole et al. (2019) found that payloads with low fitness cost can be confined if dispersal is rare [21]. However, we found that escape is highly likely for low payload fitness costs, even for a dispersal rate on the order of 10^−4^ per day.

### 5.3 Toxin-antidote recessive embryo (TARE)

The simulations of the TARE drive were extended to six years post-release as many simulations failed to reach equilibrium in the non-control region within the original four-year time period. TARE was the only gene drive we considered for which this was observed. The delayed dynamics of these simulations is likely due to proximity to a bistable equilibrium, as spatial confinement of a TARE drive requires a minimum fitness cost [15], and reduced selection in the non-control region because of the recessive fitness cost of the broken allele [43]. All metrics that we report for TARE performance are based on the payload frequency six years post-release. Fig. 6 shows the fitness profile of the TARE drive. In our modeling framework, we find that spatial confinement required a fitness cost satisfying *c* > 0.09. Threshold behavior occurs in a TARE drive when the payload fitness cost exceeds the reduction in offspring number due to the broken allele [15]. The fitness profile of a TARE drive is narrow: *c* ∈ [0.10, 0.24]. Near the edges of the fitness profile, the failure probability climbs sharply regardless of mode of failure.

**Figure 6:**
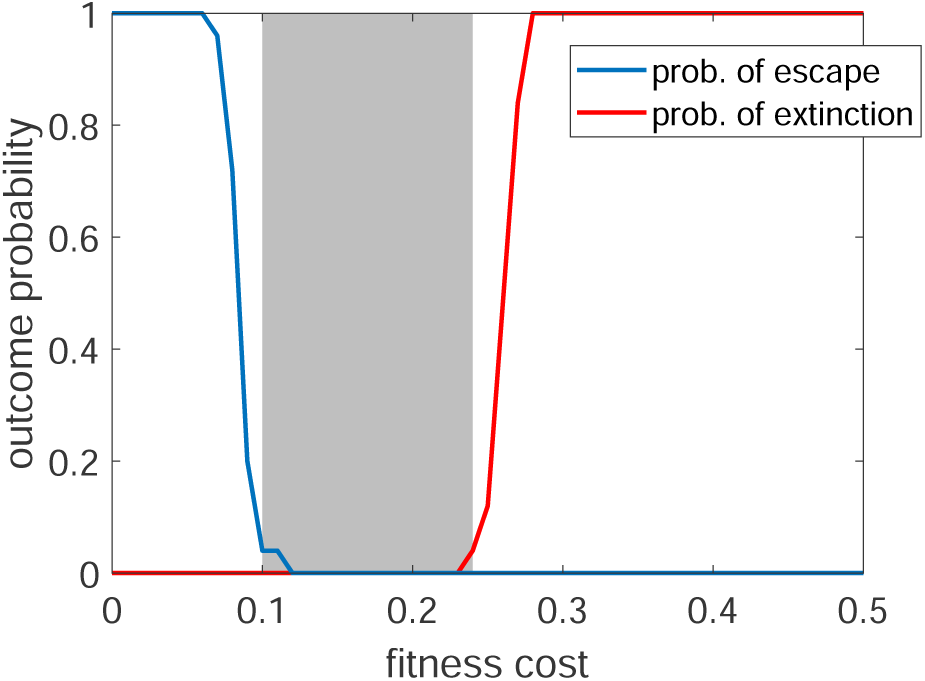
The failure probability for Toxin-Antidote Recessive Embryo (TARE). Failure is classified as either failure in persistence (extinction; red line) or failure in confinement (escape; blue line). The shaded region denotes the fitness profile of the gene drive, or the range of fitness cost values for which the failure probability does not exceed 10%, which is *c* ∈ [0.10, 0.24]. Dispersal parameters are fixed at their nominal values: the daily probability of short- and long-distance dispersal is approximately 13% and 0.04%, respectively (see Table 1). Each data point is the average of 100 stochastic simulations.

Most TARE drive simulations had a payload frequency of 80-90% in the control region, with moderate variance. The standard deviation of the payload frequency in the control region was σ̂ = 12.6% (σ̂^2^ = 1.59%; see Fig. 7). The equilibrium frequency of the payload allele decreases with increasing fitness cost as drive/wild-type heterozygotes have a higher fitness than drive homozygotes [14, 15]. Because some fitness cost is required for confinement, there is an upper limit to the equilibrium frequency that can be achieved by a TARE payload. Extinction occurred in 2.1% of simulations and was associated with high fitness cost (*c* > 0.21) and low short-distance dispersal (below 4% per day; see Fig. SM1).

**Figure 7:**
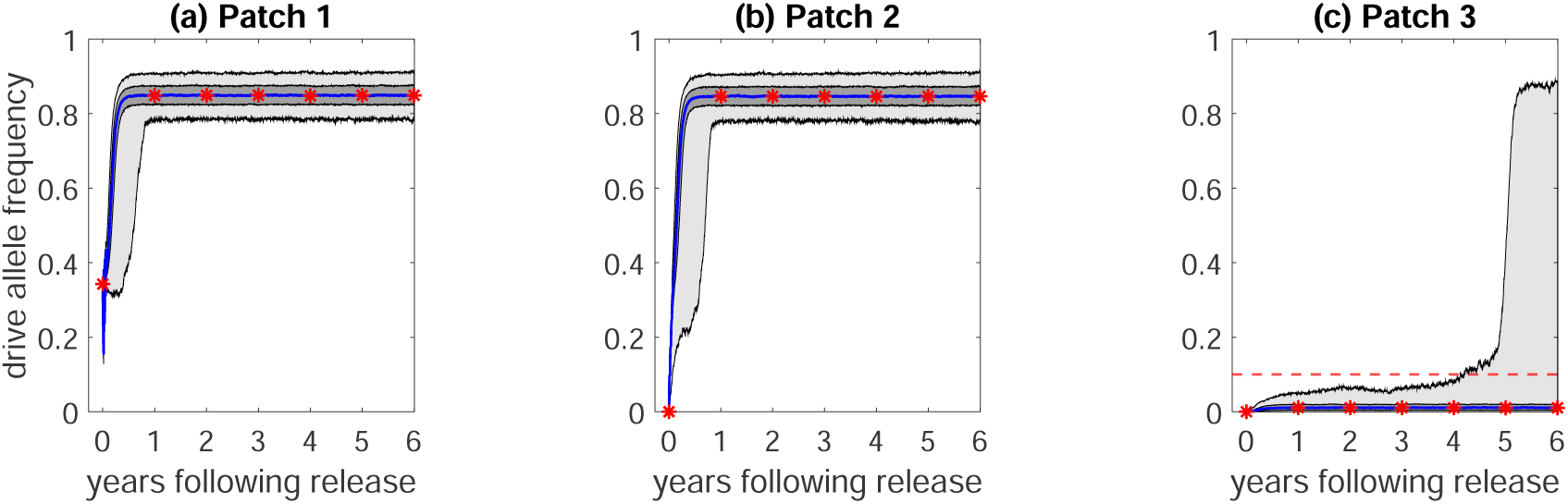
Stochastic simulations of a TARE drive in the three-patch system. The simulation time was extended to six years to allow the payload to reach numerical equilibrium in the non-control region. The middle 50th and 95th percentiles of the payload frequency are given by the light and dark gray regions, respectively. The median trajectory is denoted by the blue line with red asterisks. The plots show payload frequencies in **(a)** patch 1, **(b)** patch 2, and **(c)** patch 3. These computations are based on 1000 stochastic simulations. Note the change in *y*-axis in (c); gene drive escape occurs if the payload frequency exceeds 10%, denoted by the broken red line.

TARE drive dynamics in the control region primarily depend on payload fitness cost. The first-order sensitivity indices show that nearly 30% of allele frequency variance is due to changes in the fitness cost, while short-distance dispersal only accounted for 7%. Over 60% of payload variance is due to interactions between fitness cost and short-distance dispersal, as opposed to either parameter individually (see Table SM3). That is, the contribution of fitness cost strongly depends on short-distance dispersal and vice versa. This relationship is illustrated by the Monte Carlo simulations: no extinctions occurred in simulations with higher fitness costs when short-distance dispersal rates exceeded 10% per day. Thus, a TARE drive with a deleterious payload can spread successfully if target organism dispersal over short distances is sufficiently high.

Escape was more common than extinction, indicated by higher payload variance in the non-control region. The standard deviation of the payload frequency in the non-control region was σ̂ = 15.48% (σ̂^2^ = 2.4%; see Fig. 7(c)). Drive introgression in the non-control region is more common for several reasons. All progeny resulting from a mating between dispersing drive-bearing and wild-type organisms are viable. Consequently, payload fitness cost is the predominant factor limiting gene drive spread when frequencies are low, as the only non-viable genotype (broken allele homozygotes, i.e. ββ) rarely occurs. Furthermore, recessive fitness costs facilitate easier spread due to weaker selection [43]. These attributes contribute to a higher probability of escape, which occurred in 3.4% of the Monte Carlo simulations. Escape occurred for simulations with low fitness costs (*c* < 0.15) and long-distance dispersal rates exceeding 0.05% per day (see Fig. SM2(c)). While fitness cost was still the most influential individual source of uncertainty in payload frequency within the non-control region (16%), most of the variance was due to higher-order interactions (see Table SM2). Gene drive frequency is highly sensitive to changes in payload fitness cost when long-distance dispersal rates are relatively high.

Note that we only track the payload frequency in Figure 7(c) and not the frequency of the broken allele, which can easily achieve frequencies exceeding 10%. While the broken alleles are not themselves capable of propagating within a population, they are a byproduct of toxin-antidote systems that disrupt haplosufficient targets [55, 57].

### 5.4 Toxin-antidote dominant embryo (TADE)

TADE is similar to TARE except the broken allele formed by drive activity is dominant lethal. TADE drive confinement also requires a minimum fitness cost [14], as shown in Fig. 8. A dominant fitness effect places more selective pressure against payload spread and increases the size of the fitness profile to *c* ∈ [0.11, 0.36]. Comparing the fitness profile of TADE to that of TARE (*c* ∈ [0.10, 0.24]), we see that the lower bound of the fitness profile changes little while the upper limit increases dramatically. One benefit of TADE over TARE then is that the former is more capable of carrying costly payloads.

**Figure 8:**
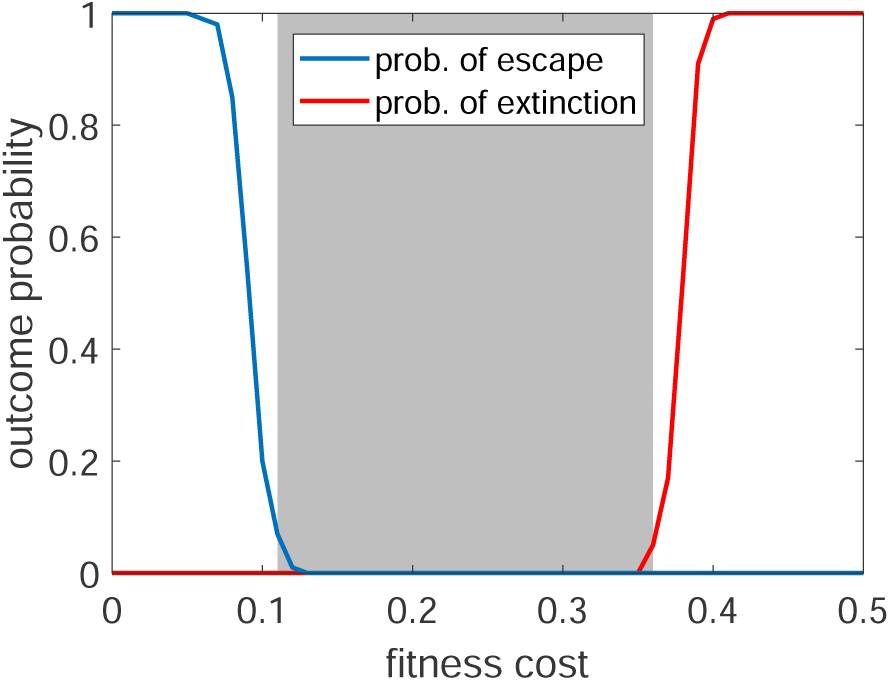
The failure probability for a Toxin-Antidote Dominant Embryo (TADE) drive. Failure is classified as either failure in persistence (extinction; red line) or failure in confinement (escape; blue line). The shaded region denotes the fitness profile of the gene drive, or the range of fitness cost values for which the failure probability does not exceed 10%, which is *c* ∈ [0.11, 0.36]. The allele frequency being measured is the payload component of the gene drive and not the broken allele. Dispersal parameters are fixed at their nominal values: the daily probability of short- and long-distance dispersal is approximately 13% and 0.04%, respectively (see Table 1). Each data point is the average of 100 stochastic simulations.

The TADE drive consistently achieved near-fixation within the control region. Equilibrium frequencies were insensitive to changes to dispersal parameters or payload fitness cost. As a result, payload variance within the control region was small: the standard deviation was σ̂ = 4.6% (σ̂^2^ = 0.2%; see Figs. 9(a)-(b)). Extinction events were exceptionally rare and occurred in only 0.5% of the Monte Carlo simulations. The simulations producing extinction tended to have relatively low short-distance dispersal, high fitness costs (*c* > 0.34), and high long-distance dispersal. Interestingly, the payload frequency in the control region is significantly affected by the long-distance dispersal rate (see Table SM2). However, given the small variance in payload frequency, arriving wild-type organisms are negligible to equilibrium dynamics.

**Figure 9:**
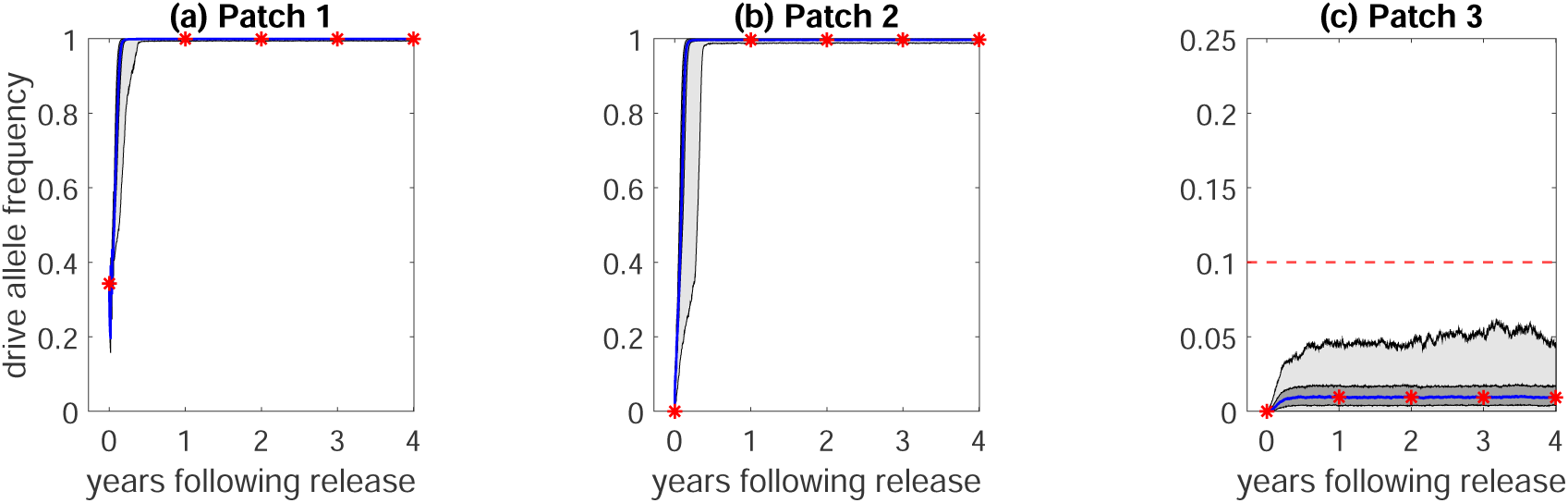
Stochastic simulations of a TADE drive in the three-patch system. The middle 50th and 95th percentiles of the payload frequency are given by the light and dark gray regions, respectively. The median trajectory is denoted by the blue line with red asterisks. The plots show payload frequencies in **(a)** patch 1, **(b)** patch 2, and **(c)** patch 3. These computations are based on 1000 stochastic simulations. Note the change in *y*-axis in (c); gene drive escape occurs if the payload frequency exceeds 10%, denoted by the broken red line.

The TADE drive showed moderate variance in the non-control region. The standard deviation of the payload frequency was σ̂ = 7.4% (σ̂^2^ = 0.55%; see Fig. 9(c)). Escape only occurred in 2.1% of the Monte Carlo simulations, and tended to occur for simulations with long-distance dispersal rates exceeding 7 × 10^−4^ or 0.07% per day. Drive escape occurred even for high payload fitness costs. The first-order indices reveal that 15.7% and 5.6% of payload frequency variance in the non-control region was due to fitness cost and long-distance dispersal, respectively. Most payload variance was due to higher-order interactions.

Our simulations show that a TADE drive rapidly spreads to near-fixation within a target population, and that this outcome is robust to changes to dispersal and payload fitness cost. However, this means that drive reversal could be practically difficult. While rapid introgression is desirable for timely modification of wild populations, this also increases the number of wild-type organisms that need to be released to reduce the payload frequency to sub-threshold levels.

## 6 Discussion

In this study, we investigated the ability of threshold gene drives to locally penetrate and remain confined to a control region. To do this, we constructed a stochastic, discrete-time patch model that incorporates different levels of dispersal. We computed the fitness profile for each gene drive, or the range of payload fitness costs for which the failure probability (i.e., the probability of extinction or escape) does not exceed 10% under nominal dispersal conditions. Next, we performed Monte Carlo simulations to quantify variance in payload frequency. Particular attention was devoted to conditions that produced gene drive failure: extinction (failed penetration) or escape (failed confinement). Finally, we conducted a variance-based sensitivity analysis to compute and compare the contribution of payload fitness cost and target organism dispersal to threshold drive performance. We outlined the conditions required for different threshold drives to achieve desirable spatial outcomes, and how threshold drive performance is sensitive to changes in fitness cost and target organism dispersal.

Several modeling assumptions and simplifications guided our analysis. Our spatial model was highly simplified and intended to provide interpretable results and insight into the dynamics of more complex spatial configurations. We found that gene drives with female-specific fitness costs are substantially harder to confine and thus assumed that all fitness costs affect both sexes equally. We did not consider gene drive resistance in our analysis. Resistance mechanisms depend on the gene drive and do not lend themselves to as simple a comparison as spatial outcomes. In practice, we anticipate that a gene drive release will incorporate strategies that reduce the probability of resistance, such as targeting highly conserved sequences and guide RNA multiplexing [16]. While our simulations used fixed parameter values, it would be interesting to investigate the effect of time-varying dispersal rates, or a comparison of threshold drive performance for changing dispersal conditions.

The variance-based analysis in this work depends on our choice of parameter priors. We used a uniform distribution in the main text, but in the Appendix we re-perform the analysis assuming that dispersal parameters are distributed according to a beta distribution. The distribution of fitness costs was left unchanged. The sensitivity indices computed using the beta prior are given in Tables SM6 and SM7. Changes in first-order indices were similar for TARE, tethered homing, and two-locus underdominance. The relative sensitivity to fitness cost tended to increase, while the relative sensitivity to either dispersal parameter tended to decrease. This follows intuitively from the beta distribution sampling values at the distribution periphery less often than the uniform distribution. For TADE, changes to first-order indices were less dramatic, with the largest change being a reduced relative sensitivity to fitness cost. The extent of higher-order interactions, measured by subtracting the first-order indices from the total-order indices, changed differently for the control and non-control regions. Higher-order interactions in the control region increased for two-locus underdominance and decreased for all remaining gene drives. A greater impact of higher-order interactions indicates that multiple parameters within the system must be modified to reduce outcome variance. This can mean that reducing performance variability is practically harder to achieve, as both fitness cost and organism dispersal must be managed. Higher-order interactions in the non-control region, in comparison, decreased for tethered homing and two-locus underdominance, increased for TARE, and stayed relatively the same for TADE. These findings underline the importance of adequately modeling target organism dispersal to inform risk analysis, as the sensitivity of gene drive performance can change depending on the distribution of parameter values. Parameter sensitivity of the TADE drive changed little for different parameter distributions, which could be desirable for a wild release.

We examined four metrics to compare the performance of threshold drives in population replacement: (1) the probability of drive failure (extinction or escape), (2) payload variance in the control and non-control regions, (3) the fitness profile, and (4) the sensitivity of gene drive performance to model parameters. These four metrics are inter-related within our modeling framework. For example, high payload variance could indicate a high sensitivity to model parameters. High payload variance in the control region can increase the probability of extinction [3], while high variance in the non-control region increases the probability of escape. If a threshold drive is insensitive to model parameters then it necessarily has low payload variance, depending on system stochasticity. Table 3 summarizes the results of these metrics for each gene drive considered in our analysis. Our results demonstrate a trade-off between gene drive persistence and confinement [8, 17, 23]. Two-locus underdominance and tethered homing were capable of robust confinement but had higher payload variance in the control region and were consequently more likely to go extinct. Furthermore, the payload frequency of these gene drives was more sensitive to changes in target organism dispersal than fitness cost. Variance in payload frequency was controlled by changes to single rather than multiple parameters. That is, 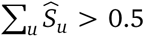 and variance in payload frequency can be reduced dramatically by modifying fitness cost independent of target organism dispersal and vice versa. Tethered homing had the widest fitness profile, so that performance was successful over a larger range of fitness costs than the other drives, even for high fitness costs. TADE had a narrower fitness profile but also performed well for costly payloads. TARE had the narrowest fitness profile. TARE and TADE had lower payload variance in the control region and thus lower extinction probabilities. However, the TARE and TADE drives had higher variance in the non-control region so that escape was more common. The toxin-antidote drives were more sensitive to changes to fitness cost than dispersal, and sensitivity was primarily characterized by higher-order interactions as 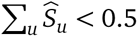. This means that the effect of modifying payload fitness cost depends on target organism dispersal and vice versa. The Monte Carlo results for TARE demonstrate that the trade-off between penetration and confinement is imperfect: the probability of both escape and extinction were higher for TARE than TADE. Part of this observation is due to a payload with a recessive fitness cost exhibiting more variation in the control and non-control regions than one with a dominant fitness cost.

**Table 3:** Results of the Monte Carlo simulations for two-locus underdominance (TLU), two-locus underdominance tethered homing (TLUTH), toxin-antidote recessive embryo (TARE), and toxin-antidote dominant embryo (TADE). The standard deviation of the payload allele frequency (σ) is computed using the drive allele frequency four years post-release (six years in the case of TARE). Estimates for the control region are obtained by averaging over patches 1 and 2. Fitness profile refers to the range of fitness costs for which the probability of threshold drive failure (extinction or escape) does not exceed 10%.

| Gene drive | Extinction prob. | Escape prob. | Control region $\hat{\sigma}$ | Non-control region $\hat{\sigma}$ | Fitness profile |
| --- | --- | --- | --- | --- | --- |
| TLU | 11% | 0% | 28.4% | 0.3% | [0.00, 0.22] |
| TLUTH | 2.7% | 0.8% | 16.6% | 1.0% | [0.04, 0.38] |
| TARE | 2.1% | 4.7% | 12.6% | 15.5% | [0.10, 0.24] |
| TADE | 0.5% | 2.1% | 7.3% | 12.0% | [0.11, 0.36] |

These findings can help guide practical deployment of a threshold drive. Two-locus underdominance and tethered homing exhibited robust confinement but moderate to high extinction probability and payload variance in the control region. Gene drives with these properties are unsuitable for large-scale replacement [70] or in regions with spatial heterogeneity [22]. High payload variance in the control region is undesirable for applications that require high payload uptake. For example, only a small fraction of females in a population of *Ae. aegypti* is capable of transmitting dengue virus at any point in time [61]. A disease-refractory payload presumably needs to achieve a high frequency in the target population to be effective. This limits the effectiveness of a TARE drive as well, as the payload frequency of a TARE drive at equilibrium has an upper limit due to a fitness cost being necessary for drive confinement. Furthermore, if human susceptibility to a disease increases due to reduced incidence, a disease-refractory payload with a fluctuating frequency in a target population could cause periodic epidemics. Robust confinement is desirable for small field releases, such as during testing [38, 45, 50], or releases into a target species that disperses over long distances (e.g., certain agricultural pests [40, 48] and mosquitoes that disperse via high-altitude winds [36]). Two-locus underdominance and tethered homing are less sensitive to changes in payload fitness cost. This is desirable in applications where fitness costs are expected to change over time, or are otherwise uncertain [42, 52]. These threshold drives were also more amenable to payloads with low fitness costs than the toxin-antidote systems. For high payload fitness costs, the tethered homing and TADE drives performed best.

The toxin-antidote drives TARE and TADE achieved better persistence in the control region but exhibited moderate to high payload variance in the non-control region and therefore higher escape probabilities compared to two-locus underdominance and tethered homing. Robust persistence is desirable when escape can be mitigated, such as when the control region is on an island (e.g., [44, 59]) or when target organism dispersal can be carefully controlled (e.g., greenhouses, crop storage facilities). The toxin-antidote drives were less sensitive to dispersal changes and thus amenable to applications for which organism dispersal between target populations fluctuates considerably, is too low to allow introgression by less efficient threshold drives (such as for sedentary target organisms, e.g. flour beetles [60]), or if there is concern of dilution by arriving wild-type organisms [32]. The TARE drive had a narrow fitness profile, indicating low tolerance for variable fitness costs, and the highest escape probability of the threshold drives we studied. These drawbacks are due to the fitness cost being recessive. Indeed, the TADE drive, which relies on a dominant fitness cost, demonstrated significantly improved performance over TARE in all metrics we investigated.

Risk management resources can be effectively allocated to monitoring variables associated with gene drive failure before, during, and after transgenic release. Extinction is comparatively rare for the toxin-antidote drives, so that monitoring resources are better allocated to tracking payload frequencies in non-control regions. In comparison, extinction was the most common source of failure for tethered homing and two-locus underdominance, so that monitoring payload frequency within the control region should take priority. Related to risk management, sensitivity to dispersal or payload fitness cost can be exploited when engineering safeguards to drive failure. The spatial outcomes of the toxin-antidote drives are sensitive to payload fitness cost, so modifications that incur high fitness costs under controlled conditions would substantially improve confinement. Examples include gene drives that restore insecticide susceptibility [6] or confer temperature-dependent fitness costs [56]. Another important aspect of risk analysis is performance uncertainty [8], which is related to drive allele variance. For example, high variance indicates a wider distribution of possible outcomes, and reducing parameter uncertainty (i.e., fixing sensitive parameters) confines gene drive performance to a narrower range. Indeed, the sensitivity analysis we performed is commonly used to quantify uncertainty.

In this work, we analyzed the performance of different threshold drives and investigated the balance between confinement and persistence using a simple spatial model. We used computer simulations to understand the variance and sensitivity of threshold drive performance to the key parameters influencing spatial outcomes: payload fitness cost and target organism dispersal. Our findings illustrate a trade-off between spatial outcomes and we demonstrate how the metrics computed here can be used to guide threshold drive deployment and risk management. These findings are useful for assessing the suitability of a threshold drive for a particular application. It remains to be explored how well this trade-off applies to other spatially-limited systems. Nonetheless, we found distinct and interesting dynamics exhibited by gene drives engineered for spatial confinement that provide greater understanding of threshold drive behavior in a wild release.

## References

[1] Akbari, O. S., Matzen, K. D., Marshall, J. M., Huang, H., Ward, C. M., and Hay, B. A., A synthetic gene drive system for local, reversible modification and suppression of insect populations, Curr Biol, 23 (2013).

[2] Alphey, L. S., Crisanti, A., Randazzo, F., and Akbari, O. S., Standardizing the definition of gene drive, Proc Natl Acad Sci U S A, 117 (2020), pp. 30864–7.

[3] Altrock, P. M., Traulsen, A., and Reed, F. A., Stability properties of underdominance in finite subdivided populations, PLoS Comput Biol, 7 (2011), p. e1002260.

[4] Altrock, P. M., Traulsen, A., Reeves, R. G., and Reed, F. A., Using underdominance to bi-stably transform local populations, J Theor Biol, 267 (2010), pp. 62–75.

[5] Anderson, M. A. E., Gonzalez, E., Ang, J. X. D., et al., *Closing the gap to effective gene drive in* Aedes aegypti *by exploiting germline regulatory elements*, Nat Commun, 14 (2023).

[6] Auradkar, A., Corder, R. M., Marshall, J. M., and Bier, E., A self-eliminating allelic-drive reverses insecticide resistance in Drosophila leaving no transgene in the population, Nat Comm, 15 (2024), p. 9961.

[7] Azzini, I., Mara, T. A., and Rosati, R., Comparison of two sets of Monte Carlo estimators of Sobol’ indices, Environ Model Softw, 144 (2021), p. 105167.

[8] Backus, G. A. and Delborne, J. A., Threshold-dependent gene drives in the wild: Spread, controllability, and ecological uncertainty, BioScience, 69 (2019), pp. 900–907.

[9] T. S. Bellows, The descriptive properties of some models for density dependence, J Anim Ecol, 50 (1981), pp. 139–56.

[10] Buchman, A., Gamez, S., Li, M., Antoshechkin, I., Li, H. H., Wang, H. W., Chen, C. H., Klein, M. J., Duchemin, J. B., Paradkar, P. N., and Akbari, O. S., *Engineered resistance to Zika virus in transgenic* Aedes aegypti *expressing a polycistronic cluster of synthetic small RNAs*, Proc Natl Acad Sci U S A, 116 (2019), pp. 3656–61.

[11] Buchman, A. B., Ivy, T., Marshall, J. M., Akbari, O. S., and Hay, B. A., *Engineered reciprocal chromosome translocations drive high threshold, reversible population replacement in* Drosophila, ACS Synth Biol, 7 (2018), pp. 1359–70.

[12] Butler, C. D. and A. L. Lloyd, How population control of pests is modulated by density dependence: The perspective of genetic biocontrol, J Theor Biol, 614 (2025).

[13] Champer, J., Champer, S. E., Kim, I. K., Clark, A. G., and Messer, P. W., Design and analysis of CRISPR-based underdominance toxin-antidote gene drives, Evol Appl, 14 (2020), pp. 1052–69.

[14] Champer, J., Kim, I. K., Champer, S. E., et al., Performance analysis of novel toxin-antidote CRISPR gene drive systems, BMC Biol, 18 (2020).

[15] Champer, J., Lee, E., Yang, E., Liu, C., Clark, A. G., and Messer, P. W., A toxin-antidote CRISPR gene drive system for regional population modification, Nat Commun, 11 (2020).

[16] Champer, J., Liu, J., Oh, S. Y., Reeves, R., Luthra, A., Oakes, N., Clark, A. G., and Messer, P. W., Reducing resistance allele formation in CRISPR gene drive, Proc Natl Acad Sci U S A, 115 (2018), pp. 5522–7.

[17] Champer, J., Zhao, J., Champer, S. E., Liu, J., and Messer, P. W., Population dynamics of underdominance gene drive systems in continuous space, ACS Synth Biol, 9 (2020), pp. 779–92.

[18] Christophers, S., Aëdes aegypti (L.) The Yellow Fever Mosquito. Its life history, bionomics and structure, Cambridge, UK: The University Press, 1960.

[19] Davis, S., Bax, N., and Grewe, P., Engineered underdominance allows efficient and economical introgression of traits into pest populations, J Theor Biol, 212 (2001), pp. 83–98.

[20] Deredec, A., Burt, A., and Godfray, H. C., The population genetics of using homing endonuclease genes in vector and pest management, Genetics, 179 (2008), pp. 2013–26.

[21] Dhole, S., Lloyd, A. L., and Gould, F., Tethered homing gene drives: A new design for spatially restricted population replacement and suppression, Evol Appl, 12 (2019), pp. 1688–702.

[22] Dhole, S., Lloyd, A. L., and Gould, F., Gene drive dynamics in natural populations: The importance of density dependence, space, and sex, Annu Rev Ecol Evol Syst, 51 (2020), pp. 505–31.

[23] Dhole, S., Vella, M. R., Lloyd, A. L., and Gould, F., Invasion and migration of spatially self-limiting gene drives: A comparative analysis, Evol Appl, 11 (2018), pp. 794–808.

[24] Edgington, M. P., Harvey-Samuel, T., and Alphey, L., Split drive killer-rescue provides a novel threshold-dependent gene drive, Sci Rep, 10 (2020), p. 20520.

[25] Edgington, M. P. and L. Alphey, Chapter 12 - modelling threshold-dependent gene drives: A case study using engineered underdominance, in Transgenic Insects: Techniques and Applications, Cabi, 2 ed., 2022, pp. 259–78.

[26] Esvelt, K. M., Smidler, A. L., Catteruccia, F., and Church, G. M., Concerning RNA-guided gene drives for the alteration of wild populations, eLife, 3 (2014), p. e03401.

[27] Gantz, V. M. and Bier, E., The mutagenic chain reaction: A method for converting heterozygous to homozygous mutations, Science, 348 (2015), pp. 442–4.

[28] Getis, A., Morrison, A. C., Gray, K., and Scott, T. W., *Characteristics of the spatial pattern of the dengue vector*, Aedes aegypti, *in Iquitos, Peru*, Am J Trop Med Hyg, 69 (2003), pp. 494–505.

[29] Grunwald, H. A., Gantz, V. M., Poplawski, G., Xu, X. S., Bier, E., and Cooper, K. L., Super-Mendelian inheritance mediated by CRISPR-Cas9 in the female mouse germline, Nature, 566 (2019), pp. 105–9.

[30] Hammond, A., Galizi, R., Kyrou, K., et al., *A CRISPR-Cas9 gene drive system targeting female reproduction in the malaria mosquito vector* Anopheles gambiae, Nat Biotechnol, 34 (2016), pp. 78–83.

[31] Harrington, L. C., Scott, T. W., Lerdthusnee, K., Coleman, R. C., Costero, A., Clark, G. G., Jones, J. J., Kitthawee, S., Kittayapong, P., Sithiprasasna, R., and Edman, J. D., *Dispersal of the dengue vector* Aedes aegypti *within and between rural communities*, Am J Trop Med Hyg, 72 (2005), pp. 209–20.

[32] Harris, K. D. and Greenbaum, G., Rescue by gene swamping as a gene drive deployment strategy, Cell Rep, 42 (2023).

[33] Hartl, D. L., A Primer of Population Genetics, Sinauer Associates, 2000.

[34] Hemme R. R., Thomas, C. L., Chadee, D. D., and Severson, D. W., *Influence of urban landscapes on population dynamics in a short-distance migrant mosquito: Evidence for the dengue vector* Aedes aegypti, PLoS Negl Trop Dis, 4 (2010), p. e634.

[35] Huang, Y., Lloyd, A. L., Legros, M., and Gould, F., Gene-drive into insect populations with age and spatial structure: A theoretical assessment, Evol Appl, 4 (2011), pp. 415–28.

[36] Huestis, D. L., Dao, A., et al., Windborne long-distance migration of malaria mosquitoes in the Sahel, Nature, 574 (2019), pp. 404–8.

[37] Kimura, M. and Weiss, G. H., The stepping stone model of population structure and the decrease of genetic correlation with distance, Genetics, 49 (1964).

[38] Knols, B. G. J., Bossin, H. C., Mukabana, W. R., and Robinson, A. S., *Transgenic mosquitoes and the fight against malaria: Managing technology push in a turbulent GMO world*, in Defining and Defeating the Intolerable Burden of Malaria III: Progress and Perspectives: Supplement to Volume 77(6) of American Journal of Tropical Medicine and Hygiene, Northbrook (IL): American Society of Tropical Medicine and Hygiene, 2007.

[39] Leftwich, P. T., Edgington, M. P., Harvey-Samuel, T., Carabajal Paladino, L. Z., Norman, V. C., and Alphey, L., Recent advances in threshold-dependent gene drives for mosquitoes, Biochem Soc Trans, 46 (2018), pp. 1203–12.

[40] Legros, M., Marshall, J. M., Macfayden, S., Hayes, K. R., Sheppard, A., and Barrett, L. G., Gene drive strategies of pest control in agricultural systems: Challenges and opportunities, Evol Appl, 14 (2021), pp. 2162–78.

[41] Li, M., Yang, T., Kandul, N. P., Bui, M., Gamez, S., Raban, R., Bennett, J., Sánchez C., H. M., Lanzaro, G. C., Schmidt, H., Lee, Y., Marshall, J. M., and Akbari, O. S., *Development of a confinable gene drive system in the human disease vector* Aedes aegypti, eLife, 9 (2020), p. e51701.

[42] Macias, V. M., Ohm, J. R., and Rasgon, J. L., Gene drive for mosquito control: Where did it come from and where are we headed?, Int J Environ Res Public Health, 14 (2017), p. 1006.

[43] Magori, K. and F. Gould, Genetically engineered underdominance for manipulation of pest populations: A deterministic model, Genetics, 172 (2006), pp. 2613–20.

[44] Marshall, J. C., Pinto, J., Charlwood, J. D., Gentile, G., Santolamazza, F., Simard, F., D. Della Torre, A., and Caccone, A., *Exploring the origin and degree of genetic isolation of* Anopheles gambiae *from the islands of São Tomé and Príncipe, potential sites for testing transgenic-based vector control*, Evol Appl, 1 (2008), pp. 631–44.

[45] Marshall, J. M. and Hay, B. A., Confinement of gene drive systems to local populations: A comparative analysis, J Theor Biol, 294 (2012), pp. 153–71.

[46] Marshall, J. M. and O. S. Akbari, Can CRISPR-based gene drive be confined in the wild? A question for molecular and population biology, ACS Chem Biol, 13 (2018), pp. 424–30.

[47] Maselko, M., Feltman, N., Upadhyay, A., Hayward, A., Das, S., Myslicki, N., Peterson, A. J., O’connor, M. B., and Smanski, M. J., Engineering multiple species-like genetic incompatibilities in insects, Nat Commun, 11 (2020).

[48] Mazzi, D. and Dorn, S., Movement of insect pests in agricultural landscapes, Ann Appl Biol, 160 (2012), pp. 97–113.

[49] Mcdonald, P. T., Population characteristics of domestic aedes aegypti (diptera: Culicidae) in villages on the kenya coast i. adult survivorship and population size, Journal of Medical Entomology, 14 (1977), p. 42–8.

[50] Nasem, Gene Drives on the Horizon: Advancing Science, Navigating Uncertainty, and Aligning Research with Public Values, Washington (DC): National Academies Press, 2016.

[51] Noble, C., Adlam, B., Church, G. M., Esvelt, K. M., and Nowak, M. A., Current CRISPR gene drive systems are likely to be highly invasive in wild populations, eLife, 7 (2018), p. e33423.

[52] North, A., Burt, A., and Godfray, H. C. J., Modelling the spatial spread of a homing endonuclease gene in a mosquito population, J Appl Ecol, 50 (2013), pp. 1216–25.

[53] North, A. R., Burt, A., and H. C. J. Godfray, Modelling the potential of genetic control of malaria mosquitoes at national scale, BMC Biol, 17 (2019).

[54] Nossent, J., Elsen, P., and Bauwens, W., Sobol’ sensitivity analysis of a complex environmental model, Environ Model Softw, 26 (2011), pp. 1515–25.

[55] Oberhofer, G., Ivy, T., and Hay, B. A., Cleave and rescue, a novel selfish genetic element and general strategy for gene drive, Proc Natl Acad Sci U S A, 116 (2019), pp. 6250–9.

[56] Oberhofer, G., Ivy, T., and Hay, B. A., Gene drive that results in addiction to a temperature-sensitive version of an essential gene triggers population collapse in Drosophila, Proc Natl Acad Sci U S A, 118 (2021), p. e2107413118.

[57] Oberhofer, G., Ivy, T., and Hay, B. A., Split versions of Cleave and Rescue selfish genetic elements for measured self limiting gene drive, PLoS Genet, 17 (2021), p. e1009385.

[58] Pianosi, F., Sarrazin, F., and Wagener, T., A Matlab toolbox for global sensitivity analysis, Environ Model Softw, 70 (2015), pp. 80–5.

[59] Pinto, J., Donnelly, M. J., Sousa, C. A., Malta-Vacas, J, Gil, V., Ferreira, C., Petrarca, V., Do Rosário, V. E., and Charlwood, J. D., *An island within an island: Genetic differentiation of* Anopheles gambiae *in São Tomé, West Africa, and its relevance to malaria vector control*, Heredity (Edinb), 91 (2003), pp. 407–14.

[60] Pointer, M. D., Spurgin, L. G., Vasudeva, R., Mcmullan, M., Butler, S., and Richardson, D. S., *Traits underlying experimentally evolved dispersal behavior in* Tribolium castaneum, J Insect Behav, 37 (2024), pp. 220–32.

[61] Rahayu, A., Saraswati, U., Supriyati, E., et al., *Prevalence and distribution of dengue virus in* Aedes aegypti *in Yogyakarta City before deployment of* Wolbachia *infected* Aedes aegypti, Int J Environ Res Public Health, 16 (2019), p. 1742.

[62] Reid, W., Williams, A. E., Sanchez-Vargas, I., Lin, J., Juncu, R., Olson, K. E., and Franz, A. W. E., *Assessing single-locus CRISPR/Cas9-based gene drive variants in the mosquito* Aedes aegypti *via single-generation crosses and modeling*, G3 (Bethesda, Md.), 12 (2022), p. jkac280.

[63] Roberts, A. J., Hackett, K., Coche, I., James, S. L., Littler, K., Santos, M., and Emerson, C. I., Taking stock: Is gene drive research delivering on its principles?, Gates Open Res, 8 (2024), p. 14.

[64] Saltelli, A., Annoni, P., Azzini, I., Campolongo, F., Ratto, M., and Tarantola, S., Variance based sensitivity analysis of model output. Design and estimator for the total sensitivity index, Comput Phys Commun, 181 (2010), pp. 259–70.

[65] Saltelli, A., Ratto, M., Andres, T., Campolongo, F., Cariboni, J., Gatelli, D., Saisana, M., and Tarantola, S., Global Sensitivity Analysis, Wiley, 2008.

[66] Sanchéz, H. M., Bennett, J. B., Wu, S. L., Rašić, G., Akbari, O. S., and Marshall, J. M., *Modeling confinement and reversibility of threshold-dependent gene drive systems in spatially-explicit* Aedes aegypti *populations*, BMC Biol, 18 (2020).

[67] Schaefer, T. J., Panda, P. K., and Wolford, R. W., Dengue fever. https://www.ncbi.nlm.nih.gov/books/NBK430732/. Date accessed: 14 Feb. 2025.

[68] Service, M. W., Mosquito Ecology: Field Sampling Methods, Chapman and Hall, London, 1993.

[69] Sheppard, P. M., Macdonal, W. W., Tonn, R. J., and Grab, B., The dynamics of an adult population of Aedes aegypti in relation to dengue heaemorrhagic fever in Bangkok, J Anim Ecol, 38 (1969).

[70] Sinkins, S. P. and F. Gould, Gene drive systems for insect disease vectors, Nat Rev Genet, 7 (2006), pp. 427–35.

[71] Trpis, M. and Hausermann, W., *Dispersal and other population parameters of* Aedes aegypti *in an African village and their possible significance in epidemiology of vector-borne diseases*, Am J Trop Med Hyg, 35 (1986), pp. 1263–79.

[72] Turelli, M. and Barton, N. H., *Why did the* Wolbachia *transinfection cross the road? Drift, deterministic dynamics, and disease control*, Evol Lett, 6 (2022).

[73] Who, Guidance framework for testing of genetically modified mosquitoes, Who, 2014.

[74] Xu, C., Legros, M., Gould, F., and Lloyd, A. L., *Understanding uncertainties in model-based predictions of* Aedes aegypti *population dynamics*, PLoS Negl Trop Dis, 4 (2010), p. e830.

[75] Yadav, A. K., Butler, C., Yamamoto, A., Patil, A. A., Lloyd, A. L., and Maxwell J. Scott, M. J., *CRISPR/Cas9-based split homing gene drive targeting* doublesex *for population suppression of the global fruit pest* Drosophila suzukii, Proc Natl Acad Sci U S A, 120 (2023), p. e2301525120.

[76] Yang, H. M., Macoris, M. L. G., Galvani, K. C., Andrighetti, M. T. M., and Wanderley, D. M. V., *Assessing the effects of temperature on the population of* Aedes aegypti, *the vector of dengue*, Epidemiol Infect, 137 (2009), pp. 1188–202.

[77] Zhang, X. Y., Trame, M. N., Lesko, L. J., and Schmidt, S., Sobol sensitivity analysis: A tool to guide the development and evaluation of systems pharmacology models, CPT Pharmacometrics Syst Pharmacol, 4 (2015), pp. 69–79.

