## Supplemental Material for "Spatial confinement of gene drives: Assessing risk of failure using global sensitivity analysis"

### Supplementary Material

Table SM1: Genotype relative fitness for each threshold-dependent gene drive we study. Fitness costs combine multiplicatively so, for example, the fitness of the $A\alpha B\beta C\gamma$ genotype for tethered homing is found by multiplying the fitnesses from their respective cells, i.e. $(1-0.05)^2(1-c)$. For tethered homing, we assume that the underdominance transgenes have fitness cost 0.05.

<table>
<tr><th colspan="6">Two-Locus Underdominance</th></tr>
<tr><td>Genotype</td><td>$AABB$</td><td>$A\alpha BB$, $AAB\beta$, $\alpha\alpha BB$, $AA\beta\beta$</td><td>$A\alpha B\beta$</td><td>$A\alpha\beta\beta$, $\alpha\alpha B\beta$</td><td>$\alpha\alpha\beta\beta$</td></tr>
<tr><td>Relative fitness</td><td>1</td><td>0</td><td>$(1-c)^2$</td><td>$(1-c)^3$</td><td>$(1-c)^4$</td></tr>
<tr><th colspan="6">Tethered Homing</th></tr>
<tr><td></td><td colspan="5">$A$, $B$ Loci</td></tr>
<tr><td>Genotype</td><td>$AABB$</td><td>$A\alpha BB$, $AAB\beta$, $\alpha\alpha BB$, $AA\beta\beta$</td><td>$A\alpha B\beta$</td><td>$A\alpha\beta\beta$, $\alpha\alpha B\beta$</td><td>$\alpha\alpha\beta\beta$</td></tr>
<tr><td>Relative fitness</td><td>1</td><td>0</td><td>$(1-0.05)^2$</td><td>$(1-0.05)^3$</td><td>$(1-0.05)^4$</td></tr>
<tr><td></td><td colspan="5">$C$ Locus</td></tr>
<tr><td>Genotype</td><td>$CC$</td><td colspan="2">$C\gamma$</td><td colspan="2">$\gamma\gamma$</td></tr>
<tr><td>Relative fitness</td><td>1</td><td colspan="2">$1-c$</td><td colspan="2">$(1-c)^2$</td></tr>
<tr><th colspan="6">Toxin Antidote Recessive Embryo</th></tr>
<tr><td>Genotype</td><td>$AA$, $A\beta$</td><td colspan="2">$A\alpha$, $\alpha\beta$</td><td>$\alpha\alpha$</td><td>$\beta\beta$</td></tr>
<tr><td>Relative fitness</td><td>1</td><td colspan="2">$1-c$</td><td>$(1-c)^2$</td><td>0</td></tr>
<tr><th colspan="6">Toxin Antidote Dominant Embryo</th></tr>
<tr><td>Genotype</td><td>$AA$</td><td colspan="2">$A\beta$, $\alpha\beta$, $\beta\beta$</td><td colspan="2">$\alpha\alpha$</td></tr>
<tr><td>Relative fitness</td><td>1</td><td colspan="2">0</td><td colspan="2">$(1-c)^2$</td></tr>
</table>

Table SM2: The total-order indices, computed using Eqtn. (7), for each parameter in the case of two-locus underdominance (TLU), two-locus underdominance tethered homing (TLUTH), toxin-antidote recessive embryo (TARE), and toxin-antidote dominant embryo (TADE). Indices are computed using the drive allele frequency four years post-release (six years in the case of TARE). Indices greater than 0.05 are considered significant. 95% confidence intervals for these indices are provided in Table SM5.

| Gene drive | Parameter | Patch 1 (%) | Patch 2 (%) | Patch 3 (%) |
|---|---|---|---|---|
| TLU | Fitness cost | 68.93 | 65.11 | 54.33 |
| | Short-distance dispersal | 73.38 | 75.26 | 8.18 |
| | Long-distance dispersal | 1.45 | 1.49 | 56.17 |
| TLUTH | Fitness cost | 23.78 | 9.17 | 77.01 |
| | Short-distance dispersal | 99.39 | 99.64 | 1.65 |
| | Long-distance dispersal | 1.43 | 1.40 | 40.84 |
| TARE | Fitness cost | 94.08 | 93.36 | 90.52 |
| | Short-distance dispersal | 69.15 | 69.15 | 0.12 |
| | Long-distance dispersal | 3.76 | 3.85 | 82.74 |
| TADE | Fitness cost | 99.24 | 95.38 | 94.39 |
| | Short-distance dispersal | 87.52 | 89.03 | 0.16 |
| | Long-distance dispersal | 11.20 | 8.99 | 84.38 |

Table SM3: The first-order indices subtracted from the total-order indices quantifies the extent of higher-order interactions between the model output and a parameter. Specifically, the difference in value is the proportion of output variance explained by higher-order parameter interactions. These differences are given for each parameter in the case of two-locus underdominance (TLU), two-locus underdominance tethered homing (TLUTH), toxin-antidote recessive embryo (TARE), and toxin-antidote dominant embryo (TADE). Indices are computed using the drive allele frequency four years post-release (six years in the case of TARE).

| **Gene drive** | **Parameter** | **Patch 1** | **Patch 2** | **Patch 3** |
|---|---|---|---|---|
| TLU | Fitness cost | 42.44 | 40.54 | 16.69 |
| | Short-distance dispersal | 43.42 | 40.64 | 5.13 |
| | Long-distance dispersal | 1.37 | 1.37 | 14.59 |
| TLUTH | Fitness cost | 23.21 | 8.97 | 19.10 |
| | Short-distance dispersal | 23.20 | 9.02 | 0.72 |
| | Long-distance dispersal | 1.41 | 1.29 | 19.06 |
| TARE | Fitness cost | 63.51 | 62.90 | 73.23 |
| | Short-distance dispersal | 63.43 | 62.83 | 0.12 |
| | Long-distance dispersal | 3.63 | 3.62 | 73.27 |
| TADE | Fitness cost | 87.61 | 85.14 | 78.73 |
| | Short-distance dispersal | 86.81 | 84.65 | 0.16 |
| | Long-distance dispersal | 11.13 | 8.75 | 78.78 |

Table SM4: 95% confidence intervals for the first-order indices in Table 2. 1000 bootstrapped samples of the sample matrices used to compute Eqtn. (6) were taken to obtain the confidence intervals reported below.

| **Gene drive** | **Parameter** | **Patch 1 (%)** | **Patch 2 (%)** | **Patch 3 (%)** |
|---|---|---|---|---|
| TLU | Fitness cost | (25.75, 27.19) | (23.86, 25.25) | (37.09, 38.23) |
| | Short-distance dispersal | (29.96, 31.74) | (33.74, 35.53) | (2.89, 3.21) |
| | Long-distance dispersal | (0.08, 0.09) | (0.11, 0.13) | (40.96, 42.17) |
| TLUTH | Fitness cost | (0.27, 0.92) | (0.11, 0.31) | (57.10, 58.64) |
| | Short-distance dispersal | (74.62, 77.68) | (89.77, 91.42) | (0.87, 1.00) |
| | Long-distance dispersal | (0.02, 0.03) | (0.10, 0.12) | (21.26, 22.33) |
| TARE | Fitness cost | (28.82, 32.42) | (28.73, 32.31) | (15.03, 19.81) |
| | Short-distance dispersal | (4.62, 7.04) | (5.22, 7.64) | (0.00, 0.01) |
| | Long-distance dispersal | (0.12, 0.15) | (0.21, 0.26) | (7.93, 11.18) |
| TADE | Fitness cost | (7.03, 16.79) | (6.53, 14.17) | (12.79, 18.40) |
| | Short-distance dispersal | (0.00, 2.18) | (2.17, 6.74) | (0.00, 0.00) |
| | Long-distance dispersal | (0.06, 0.09) | (0.21, 0.28) | (4.22, 7.17) |

Table SM5: 95% confidence intervals for the first-order indices in Table SM2. 1000 bootstrapped samples of the sample matrices used to compute Eqtn. (7) were taken to obtain the confidence intervals reported below.

| **Gene drive** | **Parameter** | **Patch 1 (%)** | **Patch 2 (%)** | **Patch 3 (%)** |
|---|---|---|---|---|
| TLU | Fitness cost | (68.03, 69.83) | (64.16, 66.01) | (53.73, 54.99) |
| | Short-distance dispersal | (72.62, 74.08) | (74.57, 75.95) | (7.93, 8.46) |
| | Long-distance dispersal | (1.28, 1.63) | (1.33, 1.66) | (55.53, 56.82) |
| TLUTH | Fitness cost | (22.30, 25.36) | (8.40, 10.07) | (76.46, 77.55) |
| | Short-distance dispersal | (99.03, 99.70) | (99.52, 99.75) | 1.55, 1.76) |
| | Long-distance dispersal | (1.06, 1.83) | (1.08, 1.75) | (40.08, 41.64) |
| TARE | Fitness cost | (92.73, 95.19) | (92.02, 94.45) | (88.82, 92.06) |
| | Short-distance dispersal | (67.24, 70.97) | (67.26, 70.95) | (0.03, 0.24) |
| | Long-distance dispersal | (3.06, 4.52) | (3.15, 4.61) | (80.24, 84.99) |
| TADE | Fitness cost | (97.76, 99.98) | (93.00, 97.63) | (92.82, 95.78) |
| | Short-distance dispersal | (82.21, 92.15) | (84.78, 92.69) | (0.03, 0.31) |
| | Long-distance dispersal | (7.65, 15.33) | (6.49, 11.91) | (81.62, 87.23) |

Table SM6: First-order indices computed using a beta distribution (distribution parameters: shape = scale = 5) scaled to the dispersal parameter range of $[10^{-6},\ 10^{-3}]$ and centered at the median. Sensitivity indices are computed for two-locus underdominance (TLU), two-locus underdominance tethered homing (TLUTH), toxin-antidote recessive embryo (TARE), and toxin-antidote dominant embryo (TADE). The indices for TARE were computed using the drive allele frequency six years post-release; all other drive indices are computed using the payload frequency four years post-release.

| **Gene drive** | **Parameter** | **Patch 1** | **Patch 2** | **Patch 3** |
|---|---|---|---|---|
| TLU | Fitness cost | 40.20 | 40.52 | 66.48 |
| | Short-distance dispersal | 8.17 | 8.39 | 0.25 |
| | Long-distance dispersal | 0.12 | 0.16 | 25.72 |
| TLUTH | Fitness cost | 56.44 | 58.28 | 84.38 |
| | Short-distance dispersal | 5.98 | 1.27 | 0.00 |
| | Long-distance dispersal | 30.64 | 35.09 | 8.24 |
| TARE | Fitness cost | 40.00 | 40.06 | 8.85 |
| | Short-distance dispersal | 3.44 | 3.75 | 0.00 |
| | Long-distance dispersal | 0.17 | 0.30 | 6.83 |
| TADE | Fitness cost | 4.03 | 4.07 | 13.80 |
| | Short-distance dispersal | 1.81 | 2.00 | 0.00 |
| | Long-distance dispersal | 0.04 | 0.19 | 8.91 |

Table SM7: Total-order indices computed using a beta distribution (distribution parameters: shape = scale = 5) scaled to the dispersal parameter range of $[10^{-6},\ 10^{-3}]$ and centered at the median. Sensitivity indices are computed for two-locus underdominance (TLU), two-locus underdominance tethered homing (TLUTH), toxin-antidote recessive embryo (TARE), and toxin-antidote dominant embryo (TADE). The indices for TARE were computed using the drive allele frequency six years post-release; all other drive indices are computed using the payload frequency four years post-release.

| **Gene drive** | **Parameter** | **Patch 1** | **Patch 2** | **Patch 3** |
|---|---|---|---|---|
| TLU | Fitness cost | 91.67 | 91.42 | 74.00 |
| | Short-distance dispersal | 59.95 | 59.59 | 1.77 |
| | Long-distance dispersal | 2.37 | 2.39 | 32.00 |
| TLUTH | Fitness cost | 62.53 | 63.65 | 91.76 |
| | Short-distance dispersal | 7.34 | 1.56 | 0.00 |
| | Long-distance dispersal | 36.33 | 40.40 | 15.62 |
| TARE | Fitness cost | 96.44 | 96.01 | 93.15 |
| | Short-distance dispersal | 59.60 | 59.44 | 0.63 |
| | Long-distance dispersal | 3.31 | 3.41 | 91.12 |
| TADE | Fitness cost | 98.06 | 97.68 | 91.16 |
| | Short-distance dispersal | 96.07 | 95.84 | 0.08 |
| | Long-distance dispersal | 9.13 | 9.24 | 86.08 |

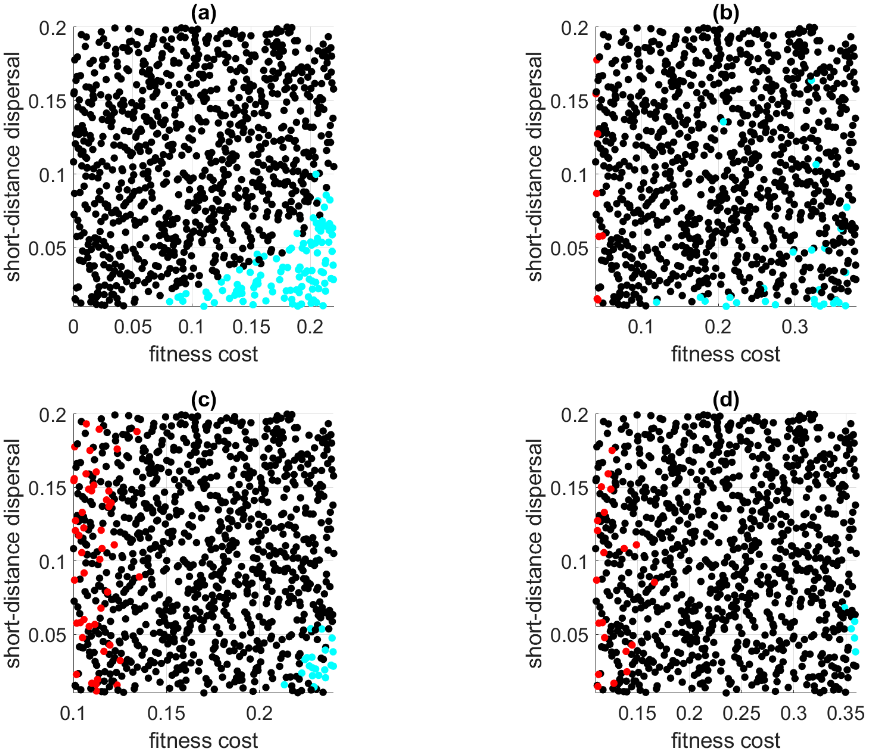


Figure SM1: Parameter combinations for the Monte Carlo simulations for **(a)** underdominance, **(b)** tethered homing, **(c)** TARE, and **(d)** TADE. Short-distance dispersal is plotted against fitness cost. Note that the $x$-axis (payload fitness cost range) differs between plots as it spans the fitness profile of each gene drive. Black dots represent individual simulations; teal dots represent parameter combinations that produced extinction, while red dots represent parameter combinations that produced escape.

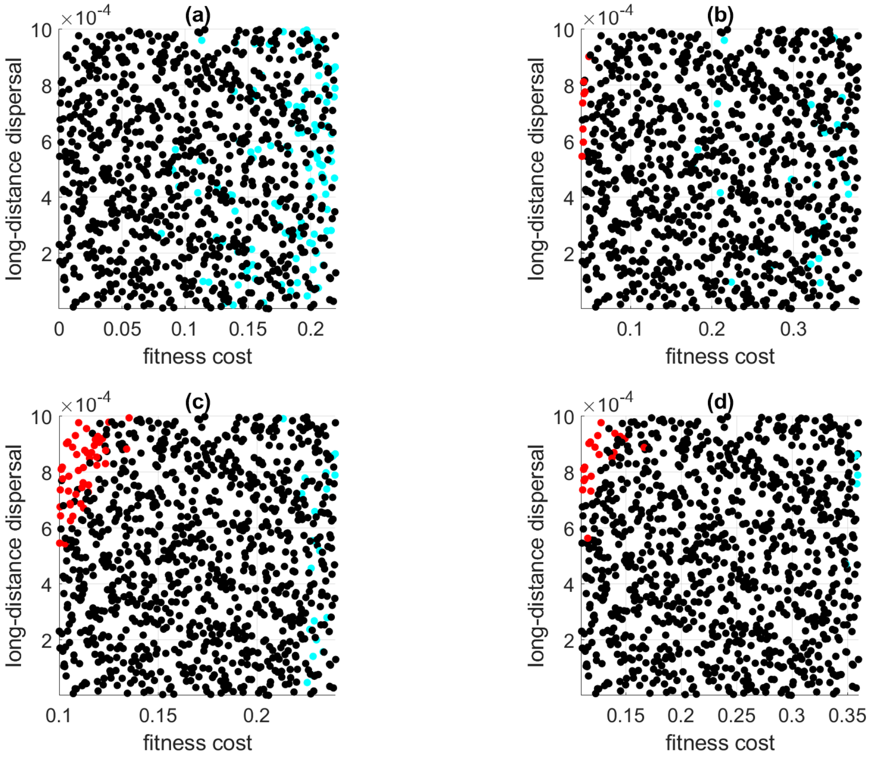


Figure SM2: Parameter combinations for the Monte Carlo simulations for **(a)** underdominance, **(b)** tethered homing, **(c)** TARE, and **(d)** TADE. Long-distance dispersal is plotted against fitness cost. Note that the $x$-axis (payload fitness cost range) differs between plots as it spans the fitness profile of each gene drive. Black dots represent individual simulations; teal dots represent parameter combinations that produced extinction, while red dots represent parameter combinations that produced escape.

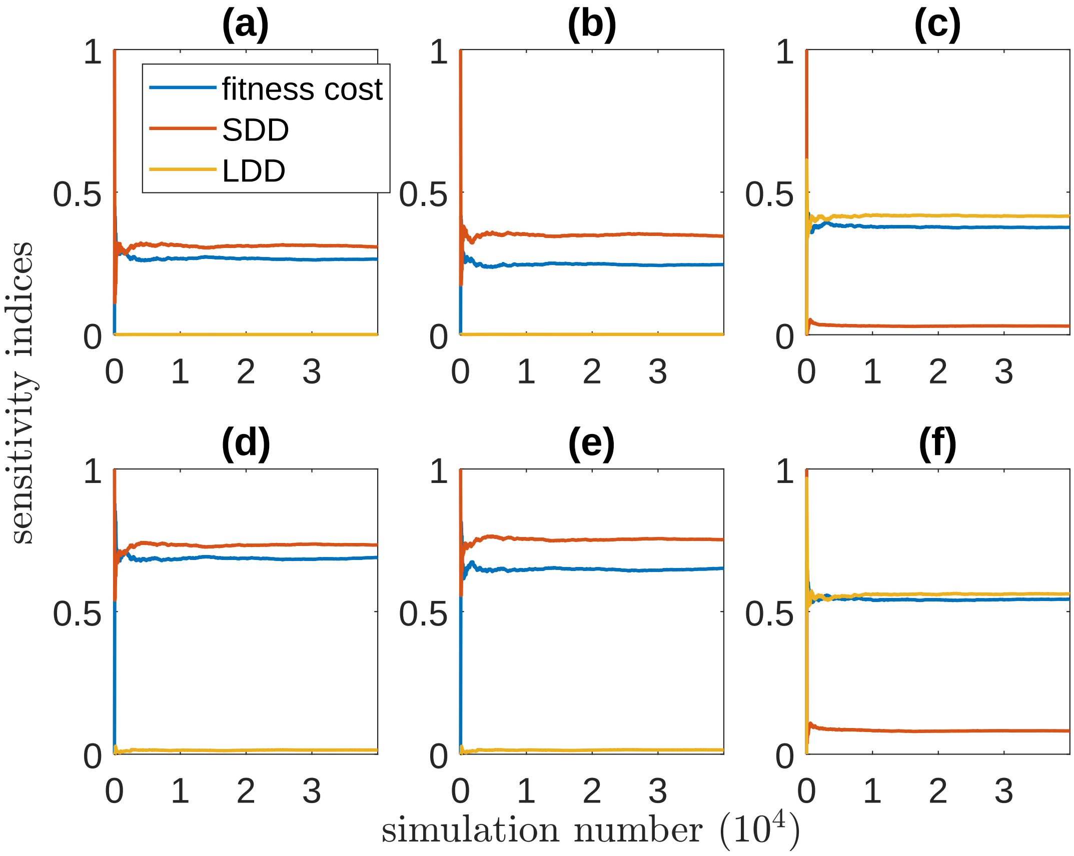


Figure SM3: Convergence of the variance-based sensitivity indices computed using Eqtns. (6) and (7) for two-locus underdominance. Each plot shows the indices for payload fitness cost, short-distance dispersal (SDD), and long-distance dispersal (LDD). We use $N = 4 \times 10^4$ simulations. The first column [**(a)** and **(d)**] corresponds to the sensitivity indices for patch 1, and similarly for the second column [**(b)** and **(e)**] and patch 2, and the third column [**(c)** and **(f)**] and patch 3. The first row shows the convergence of first-order indices while the bottom row shows convergence for the total-order indices.

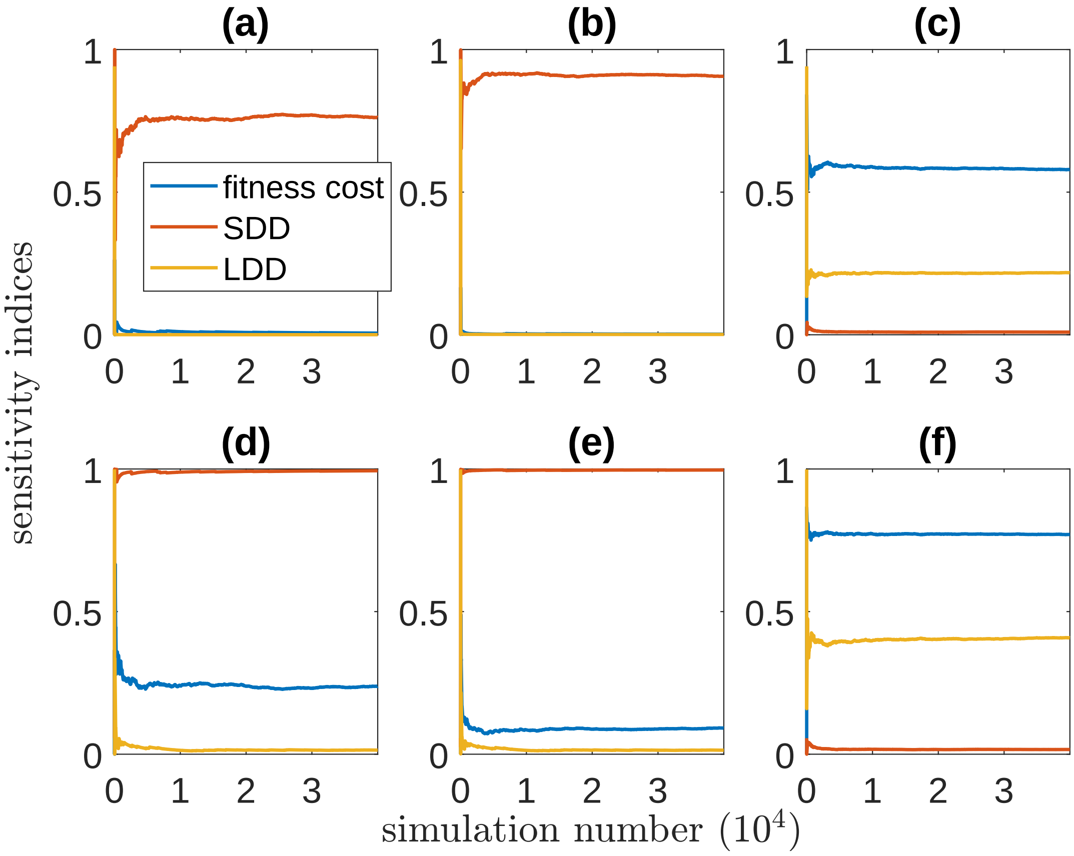


Figure SM4: Convergence of the variance-based sensitivity indices computed using Eqtns. (6) and (7) for tethered homing. Each plot shows the indices for payload fitness cost, short-distance dispersal (SDD), and long-distance dispersal (LDD). We use $N = 4 \times 10^4$ simulations. The first column [**(a)** and **(d)**] corresponds to the sensitivity indices for patch 1, and similarly for the second column [**(b)** and **(e)**] and patch 2, and the third column [**(c)** and **(f)**] and patch 3. The first row shows the convergence of first-order indices while the bottom row shows convergence for the total-order indices.

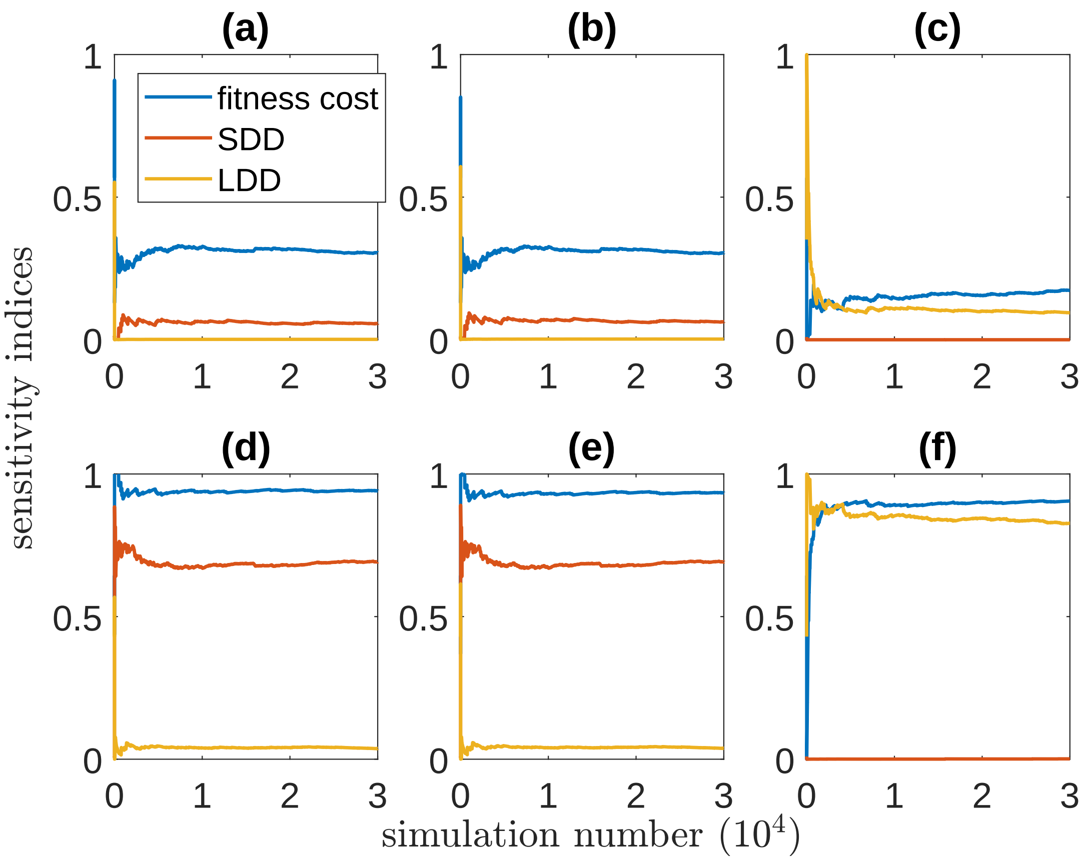


Figure SM5: Convergence of the variance-based sensitivity indices computed using Eqtns. (6) and (7) for TARE. Each plot shows the indices for payload fitness cost, short-distance dispersal (SDD), and long-distance dispersal (LDD). We use $N = 3\times10^4$ simulations. The first column [**(a)** and **(d)**] corresponds to the sensitivity indices for patch 1, and similarly for the second column [**(b)** and **(e)**] and patch 2, and the third column [**(c)** and **(f)**] and patch 3. The first row shows the convergence of first-order indices while the bottom row shows convergence for the total-order indices.

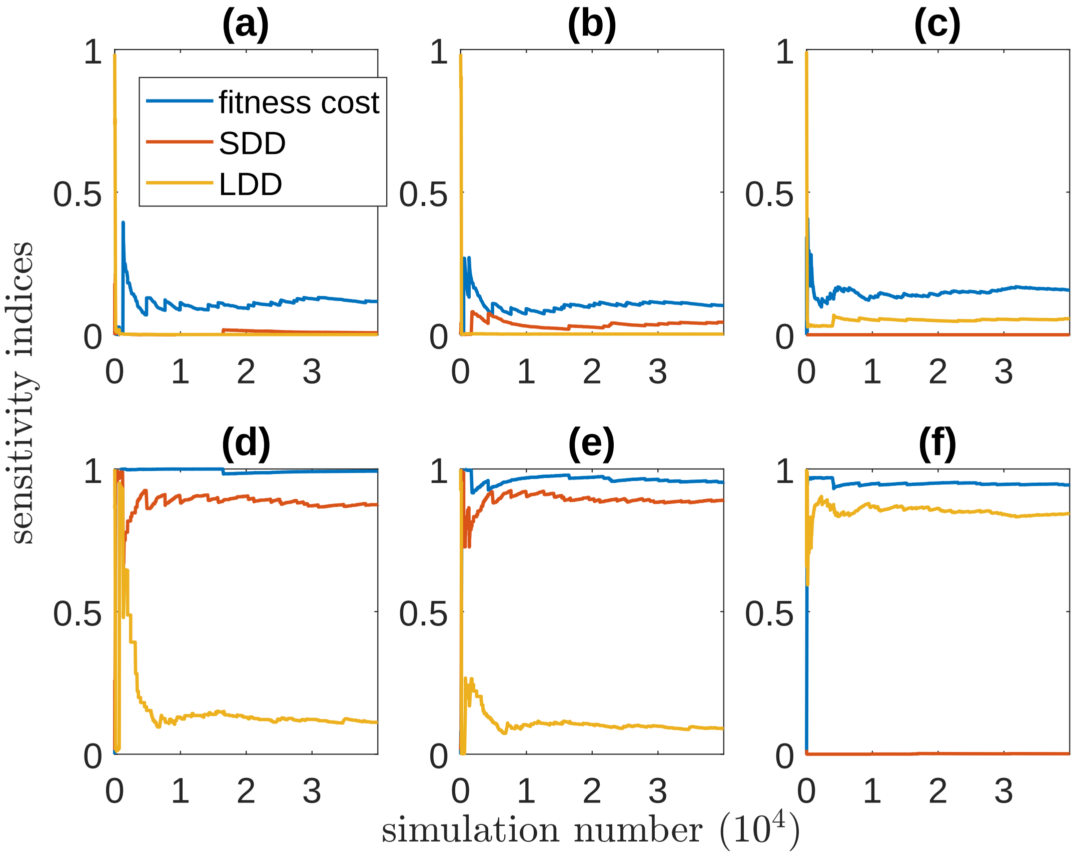


Figure SM6: Convergence of the variance-based sensitivity indices computed using Eqtns. (6) and (7) for TADE. Each plot shows the indices for payload fitness cost, short-distance dispersal (SDD), and long-distance dispersal (LDD). We use $N = 4\times10^4$ simulations. The first column [**(a)** and **(d)**] corresponds to the sensitivity indices for patch 1, and similarly for the second column [**(b)** and **(e)**] and patch 2, and the third column [**(c)** and **(f)**] and patch 3. The first row shows the convergence of first-order indices while the bottom row shows convergence for the total-order indices.
